# Changes in lipid metabolism track with the progression of neurofibrillary pathology in tauopathies

**DOI:** 10.1101/2023.09.05.556321

**Authors:** Dominika Olešová, Dana Dobešová, Petra Majerová, Radana Brumarová, Aleš Kvasnička, Štěpán Kouřil, Eva Stevens, Jozef Hanes, Ľubica Fialová, Alena Michalicová, Juraj Piešťanský, David Friedecký, Andrej Kováč

## Abstract

**Background:** Abnormal aggregation of tau protein that leads to brain inclusions is a common feature of neurodegenerative disorders called tauopathies. Recent evidence suggests the involvement of lipid metabolic deregulations in the pathogenesis of tauopathies. However, the role of tau protein in the regulation of lipid metabolism is much less characterized and not well understood.

**Methods:** We used a transgenic rat model for tauopathy to reveal metabolic alterations induced by neurofibrillary pathology. Transgenic rats express a tau fragment truncated at the N-and C-terminals. For phenotypic profiling, we performed targeted metabolomic and lipidomic analysis of brain tissue, CSF, and plasma, based on the LC-MS platform. To monitor disease progression, we employed samples from transgenic and control rats aged 4, 6, 8, 10, 12, and 14 months. To study neuron-glia interplay in lipidome changes induced by pathological tau we used well well-established multicomponent cell model system. Univariate and multivariate statistical approaches were used for data evaluation.

**Results:** We showed that tau has an important role in the deregulation of lipid metabolism. In the lipidomic study, pathological tau was associated with higher production of lipids participating in protein fibrillization, membrane reorganization, and inflammation. Interestingly, significant changes have been found in the early stages of tauopathy before the formation of high-molecular-weight tau aggregates and neurofibrillary pathology. Increased secretion of pathological tau protein *in vivo* and *in vitro* induced upregulated production of phospholipids and sphingolipids and accumulation of lipid droplets in microglia. During the later stages of tauopathy, we found a connection between the transition of tau into an insoluble fraction and changes in brain metabolism. The results showed that dysregulation of lipid composition by pathological tau leads to disruption of the microenvironment and further propagation of pathology.

**Conclusion:** Our results revealed that lipid metabolism is significantly affected during different stages of tau pathology and provide new evidence that supports the contribution of pathological tau proteins in individual lipid pathways. Our data suggests that biologically active membrane lipids such as phospholipids and sphingolipids could represent new potential next-generation therapeutic targets in tauopathies.

## Background

Tauopathies, including Alzheimer’s disease (AD), are a heterogeneous group of progressive neurodegenerative disorders characterized by severe cognitive impairment, memory loss, and aberrant behavioral patterns [1,2]. The neuropathological hallmarks of tauopathies include cerebral atrophy, hyperphosphorylation and aggregation of tau filaments into intracellular neurofibrillary tangles (NFTs), and chronic neuroinflammation. In tauopathies, post-translational modifications (PTM) of tau lead to the loss of its physiological function and subsequent progressing assembly of tau into insoluble aggregates [3]. Although the mechanism of tau aggregation is not fully understood, several factors, such as phosphorylation, acetylation, truncation, and oxidative stress, have been suggested to be able to facilitate tau fibrillization [4].

Growing evidence suggests the critical involvement of lipid dyshomeostasis in the pathogenesis of tauopathies [5,6]. Lipids are the dominant structural component of the brain and each lipid class has specific functions. Metabolism of brain lipids is closely linked to brain energy homeostasis. Under healthy conditions, lipids are important components of the cellular membrane bilayer and participate in cellular transport, energy storage, regulation of growth, and differentiation of cells. Lipids provide a hydrophobic matrix of protein-lipid/protein-protein interactions and act as essential signaling molecules and key modulators of signal transduction. Under pathological conditions, deregulation of brain lipid composition results in a disrupted blood-brain barrier, dysfunction in endocytosis, exocytosis, and autophagocytosis, altered myelination, unbalanced energy metabolism, and enhanced inflammation [7–10].

The role of tau protein in the regulation of lipid metabolism is much less characterized and not well understood. Recent studies have shown that membrane lipids like phosphatidylcholine (PC), cholesterol, and sphingolipid interact with tau protein and actively regulate its fibrillization [6,11]. Phosphatidylglycerol (PG) synthesized in mitochondria is involved in activating protein kinase C (PKC). It is possible to assume that increased activity of PKC may be involved in tau hyperphosphorylation and tangle formation [12,13].

Lipid metabolism in the brain is also closely linked to oxidative stress and neuroinflammation. Lipid accumulation in neuroglial cells affected normal neuronal activity and induced activation of microglia cells, followed by increased expression of proinflammatory mediators such as TNF-α and IL-6 [14]. Accumulation of lipids and impaired lipid metabolism are associated with the production of lipotoxic metabolites that may further trigger the progression of neurofibrillary pathology.

Both metabolites and lipids reflect the physiological and pathological status of the organism. In this study, we compared samples of transgenic (Tg) SHR-24 rat model expressing human truncated tau protein with age-matched control SHR rats. To monitor disease progression, we analyzed brain tissue (medulla oblongata, pons), CSF and plasma samples from rats aged 4, 6, 8, 10, 12, and 14 months. For phenotypic profiling, we employed targeted metabolomic and lipidomic analysis, followed by characterization of tauopathy markers. Here, we showed that lipid changes have been initiated in the early stages of tau pathology before the formation of high-molecular-weight tau aggregates. Our results indicate that lipids not only accumulate as a result of aberrant metabolic processes caused by pathological tau protein but may also contribute to aberrant protein aggregation and progression of neurofibrillary pathology. Even our multi-omics approach reveals and quantifies specific and sensitive metabolites and lipids suitable for new therapeutic interventions and playing a role in cellular pathways for early disease detection and progression of tau pathology.

## Methods

### Chemicals

Acetonitrile, methanol, isopropanol, water, methyl-tert butyl ether and ammonium acetate (LC-MS grade) were purchased from Sigma Aldrich (St. Louis, MO, USA), internal standard mixture SPLASH® LIPIDOMIX® was obtained from Avanti Polar Lipids (Alabaster, Alabama, USA) and internal standards: Creatine-d_3_ (methyl-d_3_) from CDN isotopes (Pointe-Claire, QC, Canada), L-Leucine-5,5,5-d_3_ and Butyryl-L-carnitine-(N-methyl-d_3_) hydrochloride, both from Sigma Aldrich (St. Louis, MO, USA).

### Animals

All experiments were performed on Tg SHR-24 rats expressing human truncated tau aa151-391/3R. The model was created to study the pathophysiological effects of truncated tau protein. SHR-24 rats express a tau fragment truncated at the N- and C-terminals, containing three microtubule-binding domains and a proline-rich region on an SHR (spontaneously hypertensive rat) background. Expression of the transgene is driven by the mouse Thy1 promoter. As controls, we used age-matched non-Tg animals (SHR). Animals included in this study were 4, 6, 8, 10, 12, and 14 months old. All animals were bred in the in-house animal facility of the Institute of Neuroimmunology of the Slovak Academy of Sciences in Bratislava. All animals were housed under standard laboratory conditions with free access to water and food and were kept under diurnal lighting conditions (12 h light/dark cycles with light starting at 7 am). All experiments were performed according to the institutional animal care guidelines and following international standards (Animal Research: Reporting of In Vivo Experiments guidelines) and approved by the State Veterinary and Food Administration of the Slovak Republic (Ro-933/19-221/3a) and by the Ethics Committee of the Institute of Neuroimmunology, Slovak Academy of Sciences. Efforts were made to minimize the number of animals utilized and limit discomfort, pain, or any other suffering of the experimental animals in this study.

### Human tissue specimens

Human brain tissue samples were obtained from the Queen Square Brain Bank for Neurological disorders (London, UK). All samples were obtained with informed patient consent and approval from the local ethical committees.

### Collection of samples

For the biochemical, metabolomics, and lipidomics experiments, animals were deeply anesthetized with tiletamine-zolazepam/xylazine anesthesia (4/2 mg/kg). From every rat, we collected plasma, CSF, and brain tissue. CSF samples were collected from cisterna magna in an amount ranging from 80 to 120 µL. Anesthetized animals were fixed in a head holder, and a midline incision in the skin was made up to the head area to permit easy access to the cisterna magna. After the centrifugation (4,000xg, 10 min, 4°C) samples were immediately flash-frozen in liquid nitrogen and stored at -80°C until further use. Approximately 4 ml of blood was collected from the heart to vacuum blood collection tubes with EDTA and centrifuged (4,000xg, 10 min, 4°C). Collected plasma samples were immediately frozen in liquid nitrogen and stored at -80°C. Brain tissue was cut sagittally and divided into different anatomical regions – pons and medulla oblongata. Every brain region, the left and right hemispheres, was transferred into separate tubes, flash-frozen immediately in liquid nitrogen, and stored at -80°C.

### Sample preparation

Firstly, each tissue was homogenized using homogenizer FastPrep-24 in 300 µL of methanol spiked with an internal standard mixture SPLASH® LIPIDOMIX®(2.5 %). After that, 1 ml of methyl-tert butyl ether was added and the sample was vortexed for 10s. Then, 300 µL of water was added to form a two-phase system, vortexed, and left vibrating for 15 min at room temperature (RT). The samples were subsequently left to equilibrate for 15 min at 4°C then centrifuged at 10,000g for 10 min at 4°C. At last, 400 µL of upper phase was collected for lipid analysis, and a mixture of upper and lower phase (1:1 v/v) for analysis of metabolites. Collected extracts were left to freeze at -80°C for 20 min and freeze-dried overnight. Freeze-dried samples for lipid analysis were reconstituted in 100 µL of acetonitrile/isopropanol/water (2:2:1 v/v/v), centrifuged at 10,000xg for 10 min at RT and 80 µL of supernatant was collected and transferred into LC-MS grade glass vials. For the preparation of quality control (QC) samples, 5 µL of each supernatant was mixed to form a pooled sample. Freeze-dried samples for metabolomic analysis were reconstituted in a 100 µL mixture of water/methanol (1:4 v/v). The methanolic part of the mixture contained isotopically labeled internal standards: leucine (2.0 µM), butyryl carnitine (0.2 µM), and creatinine (2.0 µM). After centrifugation (15,000xg, 15 min, 4°C) supernatants were transferred into vials, and a QC sample was prepared the same as in the sample preparation procedure for lipid analysis. CSF and plasma samples for lipid analysis were prepared by adding 80 µL of isopropanol and 2 µL of internal standard mixture SPLASH® LIPIDOMIX® (2.5%) to 20 µL of each sample, vortexed and kept overnight at -80°C for deproteinization. After centrifugation (15,000xg, 10 min, RT) 80 µL of supernatant was collected and transferred into a glass vial which was used for the subsequent LC-MS analysis. For analysis of metabolites, CSF and plasma samples were prepared by adding 20 µL of sample to 80 µL of the methanolic solution containing selected labeled internal standards, vortexed, and kept overnight at -80°C for deproteinization. Subsequently, mixtures were vortexed and centrifuged (15,000xg, 5 min, 4°C). The supernatant was transferred into vials and analyzed. The pooled QC samples for each sample type were prepared by collecting 5 µL of each sample supernatant and pooled into one.

### LC-MS analysis

Targeted lipidomic analysis was carried out by a pseudotargeted approach adopted from[15] using a liquid chromatography-tandem mass spectrometry system consisting of ExionLC^TM^ for UHPLC and QTRAP® 6500+ for MS/MS (Sciex, USA). A reversed-phase ACQUITY BEH C8 column (2.1 x 100 mm, 1.7 µm, Waters) was used for chromatographic separation. Mobile phase A consisted of acetonitrile/water (60:40 v/v), and mobile phase B consisted of isopropanol/acetonitrile (90:10 v/v) both containing 10 mM ammonium acetate. The column temperature was maintained at 55°C and the sample manager temperature was set at 15°C. Gradient starting conditions were 32%B (0 – 1.5 min) with a gradual increase to 85%B (1.5 – 15.5 min), and a further steep rise to 97%B (15.5 – 15.6 min). 97%B was maintained until 18.0 min. followed by a re-equilibration step at 32%B (18 – 20 min). The total run time was 20 min. at a flow rate of 0.35 mL/min. Mass spectra were acquired simultaneously in both positive and negative modes in the scheduled multiple-reaction monitoring (scheduled MRM) mode. The settings for the mass spectrometer were as follows: capillary voltages were set to +5500V/-4500V, the pressure of curtain gas to 40 psi, and the source temperature was 500°C. Samples were measured in randomized order with continuous monitoring of QC samples. Confirmation of correct identification of lipids was evaluated using lipid elution pattern plots (Supplementary information 1), generated by R script and the same workflow as in Drotleff, 2020 [16]. Targeted metabolomic analysis was adopted from Yuan, 2012 [17] and performed using the same LC-MS instrumentation as in the lipidomic analysis. Identification of metabolites and optimization of their measurement parameters were based on standard compounds as was already described in our previous work [18]. The aminopropyl column (Luna 3 µm NH_2_, 2 × 100 mm, Phenomenex) was used for the chromatographic separation. Mobile phase A consisted of 20 mM ammonium acetate (pH 9.75), and acetonitrile was used as mobile phase B. The column temperature was maintained at 35°C. Gradient starting conditions were as follows: 95% - 10%B (0 – 7 min), 10% B (7 – 13 min), 95%B (13 – 17 min). The total analysis time was 17 min with a flow rate of 0.3 mL/min. Mass spectra were acquired in both positive and negative modes in the scheduled MRM mode. The settings for the mass spectrometer were as follows: capillary voltages were set to +5500V/-4500V, the pressure of curtain gas to 40 psi, and the source temperature was 400°C. The data were acquired using Analyst software (v1.6.2) and processed using MultiQuant v3.0 (both Sciex, USA) for both metabolomics and lipidomics. All samples were measured in randomized order with continuous monitoring of QC samples.

### Neurofilament-light-chain and total-tau quantification in biofluids

Concentrations of neurofilament light chain (NFL) in plasma and tau proteins (total-tau) in CSF were measured using single-molecule array (Simoa) digital ELISA, using an HD-1 Analyzer. For the analysis, Simoa™ NF-Light Advantage Kit, and Simoa™ Mouse TAU Discovery Kit were used. Briefly, frozen blinded samples were melted, centrifuged for 10 min at 25°C for 20,000xg, and supernatants were transferred into a Simoa sample plate together with calibrators for NFL and Mouse TAU kits (parts of kits). Each measurement was done in duplicate according to the manufacturer’s recommendations. Concentrations were calculated using Simoa™ HD-1 instrument software.

### Cultivation and LPS stimulation of BV2 cells

Mouse microglial BV2 cells (C57BL/6, purchased from ICLC, Modena, Italy) were cultivated in Dulbecco’s Modified Eagle’s Medium (DMEM, PAA Laboratories GmbH, Colbe, Germany) containing 10% fetal calf serum (FCS, Invitrogen, Carlsbad, California, USA), and 2 mM L-glutamine (PAA laboratories GmbH, Colbe, Germany) at 37°C and 5% CO_2_. The medium was changed twice a week. One day before treating the BV2 cells with recombinant tau protein (aa 151-391/3R), the cell culture medium was replaced with a serum-free DMEM medium with L-glutamine. To stimulate the cells, tau protein was added for a final molarity of 500 nM for 24 hours.

### Isolation and cultivation of primary rat glial culture

Rat mixed glial culture was prepared from cerebral cortices of 1–2 day old Sprague Dawley rats (n = 8/isolation). The animals were euthanized by CO_2,_ and the cerebral cortices were dissected, stripped of the meninges, and mechanically dissociated by repeated pipetting followed by passage through a 20 µm nylon mesh (BD Falcon, Franklin Lakes, USA). Cells were plated on 6-well plates pre-coated with poly-L-lysine (10 µg/mL, Sigma-Aldrich, St. Louis, MO) and cultivated in DMEM medium containing 10% FCS and 2 mM L-glutamine at 37°C, 5% CO_2_ in a water-saturated atmosphere.

### Development of a multi-component cell model system

The multi-component model was composed of rat primary microglia, astroglial cells, and human neuroblastoma cell line SH-SY5Y with inducible expression of truncated tau protein (previously described by [19]). Firstly, the expression of truncated tau protein was induced by cultivating cells in a medium without doxycycline (Sigma-Aldrich, St. Louis, MO) for three days before cell seeding into co-culture. Secondly, SH-SY5Y cells were co-cultivated with primary glial culture in 6-well plates in density 1.10^3^ cells/cm^2^ for a further 5 days in DMEM medium containing 10% FCS, 2 mM L-glutamine and N2-supplement (Life Technologies Invitrogen, Carlsbad, CA) at 37°C and 5% CO_2_. The medium was changed twice a week. 24 hours before the experiment, we cultivated the cells in a medium with 2% fatty acid-free BSA (Life Technologies Invitrogen, Carlsbad, CA).

### Isolation of lipid droplet-associated mitochondria (LDM)

The brainstem from Tg and control rats (n = 4) was harvested and washed with phosphate buffer saline (PBS, 137 mM NaCl, 2.7 mM KCl, 10 mM Na_2_HPO_4_, 2 mM KH_2_PO_4_, pH 7.4) to remove blood contamination. The brain tissue was suspended in sucrose-HEPES-EGTA buffer supplemented with 2% BSA (250 mM Sucrose, 5 mM HEPES, 2 mM EGTA, 2% fatty acid-free BSA, pH 7.2). The tissue suspension was homogenized and the homogenate was transferred into a 15 mL falcon tube. Homogenate was centrifuged in a swinging bucket rotor at 900xg for 10 min at 4°C. The post-nuclear supernatant was collected and layered with Buffer B (20 mM HEPES, 100 mM KCl, 2 mM MgCl_2_, pH 7.4). The second centrifugation was carried out in a swinging bucket rotor at 2,000xg for 40 min at 4°C to allow the fat layer to separate. The layer that contains LDs appears as a translucent band on the top of the gradient. This layer was resuspended in HEPES buffer without BSA and centrifuged at 10,400xg for 10 min at 4°C in a fixed rotor. The LDM pellet was re-suspended in mitochondrial resuspension buffer (250 mM mannitol, 5 mM HEPES, 0.5 mM EGTA, pH 7.4).

### ATP assay

The assay was performed using a Luminescent ATP detection assay kit from Promega (Wisconsin, USA). 50 µL of isolated mitochondria (LDM) were suspended in 50 µL of 20 mM Tris pH 7.5 buffer and combined with 100 μL of CellTiter-Glo® 2.0 Reagent in an ELISA standard plate. The plate was placed on an orbital shaker for 5 min., and luminescence was recorded immediately thereafter using Luminometer, Fluoroscan Ascent FL (ThermoScientific, US).

### AlamarBlue Assay

AlamarBlue™ solution was added to the cell culture in dilution 1/10. Cells were incubated at 37 °C and 5 % CO_2_ for 4 hours. 100 µl aliquots of each sample were transferred to replicate wells of a black 96-well plate. The fluorescence was measured at 590 nm, using an excitation wavelength of 530-560 nm.

### Immunocytochemistry and BODIPY staining

BV2 and primary rat glial cells were fixed for 15 min in 4% paraformaldehyde (Sigma-Aldrich, St. Louis, MO, USA) and washed with PBS. Cells were blocked for 60 min in 5% BSA in PBS. Cells were washed with PBS and incubated with primary antibody rabbit polyclonal anti-IBA-1 (FUJIFILM Wako Chemicals, USA) overnight. After washing, the cells were stained with BODIPY 493/503 (1:1000, Thermo Fisher) for 15 min at RT. Cells were mounted and examined using an LSM 710 confocal microscope (Zeiss, Jena, Germany).

### Immunohistochemistry and BODIPY staining

Animals were anesthetized by tiletamine-zolazepam/xylazine anesthesia and perfused intracardially with PBS with heparin. The brain was removed and embedded in a cryostat embedding medium (Leica, Wetzlar, Germany) and frozen above the surface of liquid nitrogen. 12-μm-thick brain sections were cut on a cryomicrotome, fixed onto poly-L-lysine coated slides, and left to dry at RT for 1 hour. Sections were fixed for 15 min in 4% paraformaldehyde and blocked for 60 min in a blocking solution (DAKO, Ontario, Canada). Sections were incubated overnight in a polyclonal rabbit anti-IBA-1 primary antibody (1:1000, Wako), polyclonal rabbit anti-GLUT3 primary antibody (1:100, Invitrogen, Massachusetts, USA), and monoclonal mouse anti-NeuN primary antibody (1:1000, Abcam, Waltham, MA, USA). After washing, the sections were incubated in goat anti-rabbit or anti/mouse AlexaFluor546/488 secondary antibody (1:2000, Invitrogen, Massachusetts, USA) for 1 hour at RT. Sections were washed with PBS and incubated with BODIPY 493/503 in PBS (1:1000, Invitrogen, Massachusetts, USA) for 15 min at RT. For tau staining, sections were incubated in monoclonal mouse anti-rat pSer202/pThr205 primary antibody (1:1000, Invitrogen, Massachusetts, USA). After washing, the sections were incubated for 1 hour in a secondary antibody goat anti-mouse AlexaFluor488 (1:1000, Invitrogen, Massachusetts, USA). Brain slices were mounted (Vector Laboratories, Burlingame, CA, USA) and examined using an LSM 710 confocal microscope. Oil Red O (Abcam, Waltham, MA, USA) was used for the staining of neutral lipids. The slides were incubated in propylene glycol for 5 min and then in heated Oil Red O solution for 10 min. The sections were differentiated in 85% propylene glycol for 1 min and washed twice in water. The brain slices were mounted in an aqueous mounting medium.

### Measurement of membrane fluidity

Membrane fluidity was measured by lipophilic pyrene probe - pyrenedecanoic acid (PDA, Abcam, Waltham, MA, USA), which undergoes dimerization after the interaction and exhibits changes in its fluorescent properties. BV2 cells were treated with tau protein (final concentration 1 um) for 24 hours. The cells were incubated with PDA/Pluronic F127 solution for 1 hour at 25°C in the dark with gentle agitation. The unincorporated PDA was removed by washing cells twice with cultivation media. The live-cell fluorescence microscopy was performed. Changes in cell membrane lipid order between the monomer gel/liquid-ordered phase (fluorescent at 430∼470 nm) and the excimer liquid phase (fluorescent at 480∼550 nm) were measured by confocal microscopy.

### Preparation of sarkosyl-insoluble PHF-tau from Tg rat brain

Purified sarkosyl-insoluble PHF-tau from Tg rat brains was isolated according to a previously published protocol [20]. For isolation of sarkosyl-insoluble tau, we used brain tissue from 4, 6, 8, 10, 12, and 14-month-old Tg animals (n= 6). Briefly, 20 mg of the medulla and pons from a Tg rat brain enriched in PHF-tau were dissected, cleaned of blood vessels and meninges, and used as a starting material. The brain tissue was homogenized on ice in 10 volumes of ice-cold SL buffer (20 mM Tris-HCl, pH 7.4, 800 mM NaCl, 1 mM EGTA, 1mM EDTA, 0.5% β-mercaptoethanol, 10% sucrose, 1x protease inhibitors Complete, EDTA-free). The homogenate was centrifuged for 20 min at 20,000xg. Solid sarkosyl (Sigma-Aldrich, St. Louis, MO, USA) was added to the supernatant to achieve a 1% concentration. The samples were stirred for 1 hour at RT. The samples were spun for 1 hour at RT in a Beckman ultracentrifuge using SW 120 Ti rotor (Beckman Coulter, Inc., Brea, CA, USA) at 100,000xg. After sarkosyl extraction and ultracentrifugation, pellets containing PHF-tau were dissolved in a 1 x SDS sample loading buffer and boiled for 5 min. Samples were determined by following western blot analysis.

### Biochemical western blot analysis

Samples were separated on 12% SDS-polyacrylamide gels and transferred to a nitrocellulose membrane in 10 mM N-cyclohexyl-3-aminopropanesulfonic acid (CAPS, pH 11, Roth, Karlsruhe, Germany). The membranes were blocked in 5% milk in Tris-buffered saline with 0.1% Tween 20 (Sigma-Aldrich, St. Louis, MO, USA) (TBS-T, 137 mM NaCl, 20 mM Tris-base, pH 7.4, 0.1% Tween 20) for 1 hour and incubated with primary antibody overnight at 4°C. As phospho-dependent anti-tau antibodies, we used: mouse monoclonal anti-rat pThr181 (1:1000, Invitrogen Life Technologies, Carlsbad, CA), mouse monoclonal anti-rat pSer202/pThr205 (AT8, 1:1000, Invitrogen Life Technologies, Carlsbad, CA), mouse monoclonal anti-rat pThr231 (1:1000, Invitrogen Life Technologies, Carlsbad, CA), mouse monoclonal anti-rat pThr181 (1:1000, Invitrogen Life Technologies, Carlsbad, CA), rabbit polyclonal anti-rat pSer199 (1:1000, Invitrogen Life Technologies, Carlsbad, CA), rabbit polyclonal anti-rat pSer262 (1:1000, Invitrogen Life Technologies, Carlsbad, CA), mouse monoclonal anti-rat DC217 (1:200, Axon Neuroscience R&D SE, Bratislava, Slovakia) and polyclonal anti-rat pThr212 (1:1000, Invitrogen Life Technologies, Carlsbad, CA). For total tau, we used a mouse monoclonal anti-rat DC25 antibody (recognizing epitope 368–376, 1:200, Axon Neuroscience R&D SE, Bratislava, Slovakia). Membranes were incubated with horseradish peroxidase (HRP)-conjugated secondary antibody in TBS-T (1:3000, Dako, Glostrup, Denmark) for 1 hour at RT. Immunoreactive proteins were detected by chemiluminescence (SuperSignal West Pico Chemiluminescent Substrate, Thermo Scientific, Pittsburgh, USA) and the signals were digitized by Image Reader LAS-3000 (FUJIFILM, Bratislava, Slovakia). The signal was semi-quantified by ImageJ software.

### Quantitative analysis of immunocytochemical and immunohistological data

Relative staining patterns and intensity of projections from immunohistochemical and immunocytochemical stainings were visualized by confocal microscopy and evaluated. ImageJ (public domain ImageJ software) was used for the evaluation and quantification. We quantified 10 slices from each sample. For semiquantitative analysis, the color pictures were converted to grayscale 8-bit TIFF file format, and regions of interest were analyzed. The grayscale 8-bit images were converted to 1-bit images, on which the number of immunolabeled structures localized in the area of interest was measured. The average intensity/pixel values of each area were then calculated, and the average intensity/pixel values representing the background intensity were subtracted from those of the immunolabeled areas.

### Data analysis

Lipidomics and metabolomics data were processed and statistically evaluated in R software (www.r-project.org, 2019, v 3.5.0) using the Metabol package[21]. At first, the quality control-based locally estimated smoothing signal correction (LOESS) was applied to each dataset. The instrumental signal stability was monitored by internal standards across experimental and QC samples. The coefficient of variation (CV), based on QC samples, was calculated, and compounds (lipids or metabolites) with a CV higher than 30% were excluded from further processing. In the case of the brain, the correction of datasets to sample weight was carried out. The data were transformed by natural logarithm (ln), and the mean centering was applied. Bonferroni correction of p-values was applied to prevent false positivity of the t-test. Univariate and multivariate statistical methods were used to evaluate and visualize the results. Cytoscape software v3.8.2 was used to create metabolic and lipid maps according to the results of statistical analyses[22]. For metabolomics data, an Enrichment pathway analysis was performed using the web-based platform MetaboAnalyst (v5.0)[23] using our in-house database, which contained up to 53 metabolic pathways (Table S1). The lipid ontology (LION) enrichment analysis web application was used for the enrichment analysis of lipidomic data [24]. Data from western blot analysis, immunochemistry, ELISA, and FACS were statistically analysed using GraphPad Prism software v.8.0.1 (Inc., La Jolla, CA, USA).

## Results

### The transition of tau into high-molecular forms progressively increases in an age-dependent manner

Pathological post-translational modifications of tau protein led to its misfolding, aggregation, and progressive accumulation in brain tissue. To monitor the development of tau pathology in an age-dependent manner, we performed an immunohistochemical staining and semi-quantitative analysis of sarkosyl insoluble protein extracts from the pons and medulla oblongata of SHR-24 Tg rats at the ages of 4 to 14 months. The results showed the age-dependent increase of insoluble pathological phospho-tau species using AT8 (pSer202/pThr205) immunostaining in pons and medulla (Fig. 1A, B). To detect soluble and sarkosyl-insoluble hyperphosphorylated forms of tau in brain extracts, we used antibodies against various phospho-epitopes: pSer202/pThr205, pThr217, pThr212, pThr181, pSer262, pThr231, and pSer199. Total tau was determined with a DC25 (epitope aa 368–376) anti-tau antibody. Biochemical results confirmed the pathological changes in the brain tau protein. There were no high molecular weight aggregates in sarkosyl-insoluble tau in pons (Fig. 1C) and medulla (Fig. 1D) from 4- and 6-month-old Tg rats. Only the aggregated truncated tau (aa 151-391/3R) was present. In 8-month-old animals, endogenous tau co-assembled with truncated tau and became a part of the sarkosyl-insoluble complexes. It is important to note that we observed differences in insoluble tau aggregates in studied brain regions, the pons was affected to a lesser extent than the medulla. In Tg rats, there was an age-dependent increase in tau phosphorylation. We found that phosphorylations at pSer202/pThr205, pThr217, pThr212, and pThr181 were significantly increased in the medulla in comparison to the pons. The highest amount of phosphorylated and aggregated tau was detected in 10- and 14-month-old animals (Fig. 1E, F). Interestingly, pThr217 was present in high molecular weight insoluble fraction (50-70kDa) in the medulla already in 8 months of age, supporting its crucial role in tau pathology development [25].

**Fig. 1.**
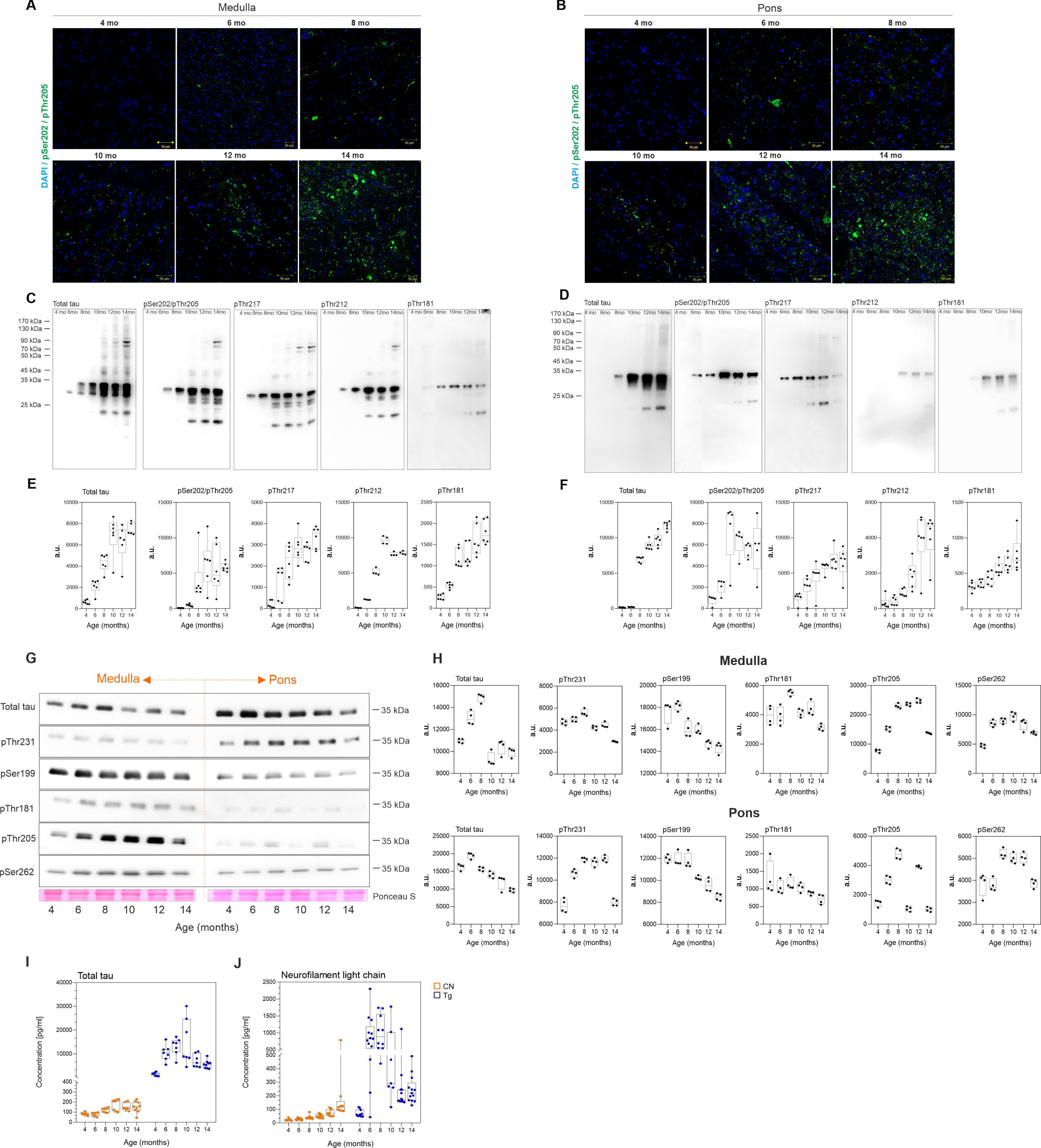
Neurofibrillary pathology progressively increased in an age-dependent manner in the brainstem of Tg rats. **A-B,** Immunohistochemical staining (DAPI-blue, pSer202/pThr205-green) demonstrated the presence of tau protein phosphorylated at pSer202/pThr205 in the medulla oblongata and pons of Tg rats. **C-F,** Progressive increase in the levels of sarkosyl-insoluble tau protein complexes in the medulla oblongata and pons of Tg rats. **G-H**, Increase in soluble hyperphosphorylated truncated tau in medulla and pons of Tg rats. Ontogenesis of soluble tau and sarkosyl-insoluble tau complexes in aging rats from 4 to 14 months (n= 6 for each age) was monitored by Western blot analysis using phosphorylation-dependent anti-tau antibodies against pThr212, pThr181, pThr217, pSer199, pThr231, pSer262 and pSer202/pThr205. Total tau was determined with a DC25 anti-tau antibody. **I-J,** The levels of biofluid markers (neurofilament and total tau) of tau pathology. Data are presented as boxplots (Min to Max).

We also analyzed soluble fractions of truncated tau protein in brain extracts from pons and medulla oblongata (Fig. 1G) where soluble tau as well as hyperphosphorylated protein were increased in 6-month-old Tg rats and then slowly decreased in time due to the development of high-molecular-weight aggregates (Fig. 1H). Significant trends suggested increased expression of pSer199, pThr205, and pSer262 up to the 6^th^ month of age.

Biofluid markers of tau pathology were measured in plasma and CSF. The CSF total-tau levels were significantly elevated in 4-month-old SHR-24 Tg rats and increased gradually up to the 10^th^ month of age. In CSF, we observed a slight decrease in tau levels in 12- and 14-month-old animals (Fig. 1I, Table S2). Decreasing levels of total tau in the CSF after 12 months correlated with the appearance of high molecular tau aggregates in brain tissue and the upregulation of soluble hyperphosphorylated tau forms. Similarly, elevated neurofilament light chain (NFL) levels were found in the plasma of 4-month-old SHR-24 Tg rats and older. NFL plasma levels remained high up to the 10^th^ month of age and started to decrease in 12-month-old animals (Fig. 1J, Table S2). Similarly, to tau protein, NFL might aggregate in brain tissue, hence its release to biofluids decreases with progressing damage.

### Brain lipidome and metabolome are significantly altered by tau pathology

To precisely investigate the impact of tau-induced neurodegeneration on brain metabolism and uncover the lipids and metabolites involved, we performed targeted lipidomic and metabolomic analysis of brain tissues, CSF, and plasma samples of Tg rats and their respective age-matched control groups at 4 to 14-months of age (Fig. 2A). Using UHPLC-MS/MS analysis we found 722/150 and 894/143 lipid species and metabolites in the medulla oblongata and pons. In CSF and plasma, we detected 325/76 and 759/129 lipids and metabolites (Fig. 2B, C). The summary and abbreviations of detected lipids and metabolites are in Table S3.

**Fig.2.**
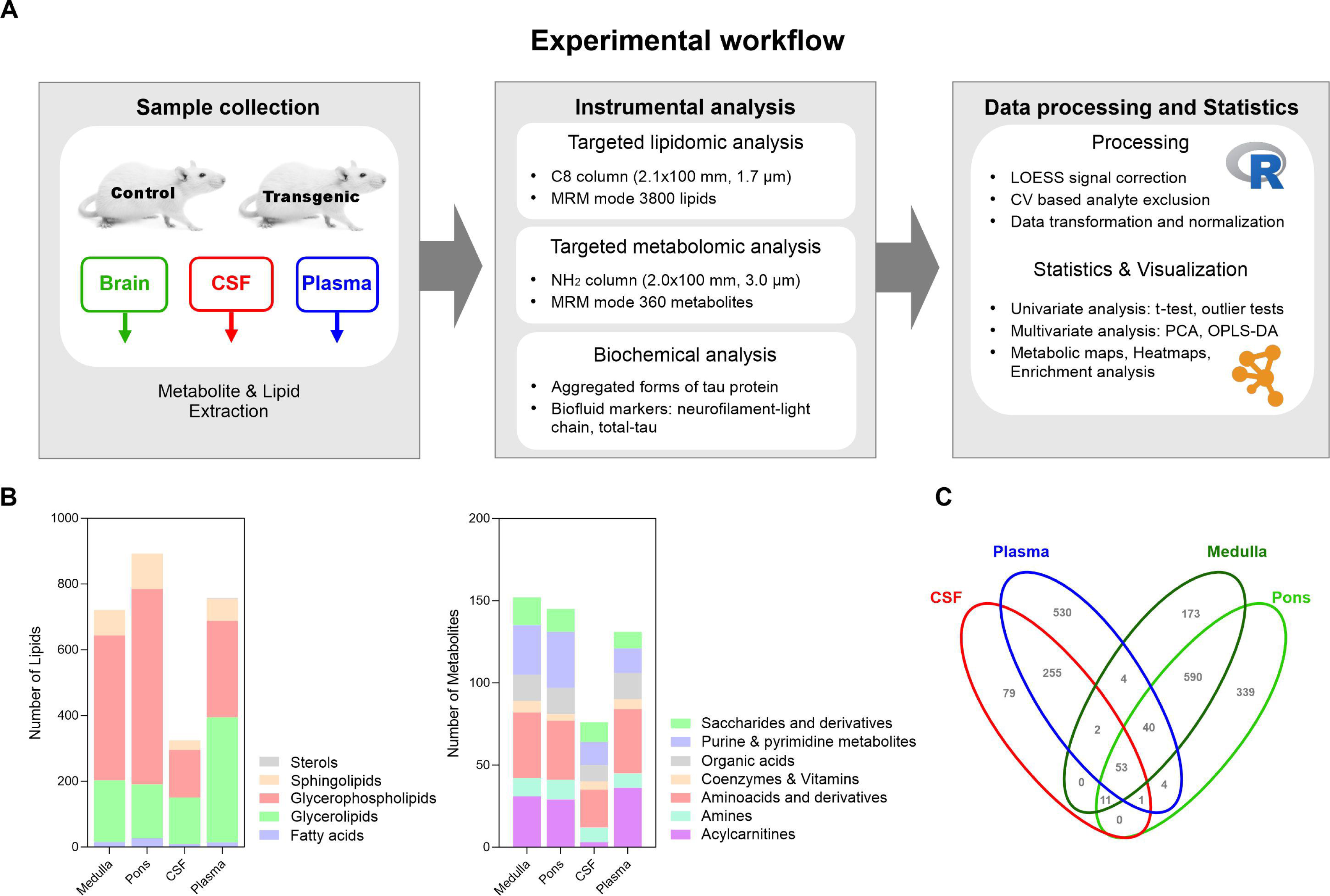
Overview of experimental approach, targeted lipidomic and metabolomic analyses, and data processing. **A** Experimental workflow started with the collection of brain tissue, CSF, and plasma samples of 4, 6, 8, 10, 12, and 14 months-old transgenic (SHR-24) and age-matched controls (SHR). Subsequently, high-coverage lipidomic and metabolomic analyses were performed together with biochemical analysis for tau pathology characterization. Two omics datasets were processed and statistically evaluated together using several approaches. **B** Overview of several identified lipids and metabolites in brain tissue, CSF, and plasma. Lipids were grouped according to the LIPID maps classification system (fatty acids, glycerolipids, glycerophospholipids, sphingolipids, and sterols). Metabolites were grouped according to their chemical structure into 7 main groups (acylcarnitines, amines, amino acids, and derivatives, coenzymes, and vitamins, organic acids, purine & pyrimidine metabolites, saccharides, and derivatives). **C** Venn diagram shows the numbers of identified lipids and metabolites in each compartment.

Our results demonstrated distinct metabolic patterns in the medulla oblongata of Tg rats as illustrated in principal component analysis (PCA), and orthogonal partial least-square discriminant analysis (OPLS-DA) calculated from joined lipidomic and metabolomic datasets (Fig. 3A). In unsupervised PCA score plot a gradual separation according to the age with baseline separation of old from young animals in both groups was present. When comparing Tg rats with controls, we observed partial separation of age-matched groups as soon as the 8^th^ month of age. Group separation was more apparent with progressing age, peaking in 10, 12, and 14-month-old animals. Supervised OPLS-DA analysis results in a clearer representation of observed trends in PCA. In pons, PCA followed partly similar trends as in medulla oblongata, although to a lesser extent (Fig. S1A). To further investigate the metabolic patterns and individual analytes contributing to the group separation, we used univariate statistics (student’s t-test corrected on multiple comparisons by Bonferroni) plotted on a metabolic network (Fig. 3B, F, Fig. S1B) and summarized as a heatmap in Table S4. In medulla oblongata, the accumulation of pathological tau peaked at 10 months of age, thus we used this age-matched group as a starting point for univariate statistics. When comparing the lipid profiles of 10-month-old Tg rats vs. controls, major changes were attributed to glycerophospholipids (GPL). A considerable increase in lysophospholipid (lysoPL) levels was observed especially in lysophosphatidylethanolamine (LPE), lysophosphatidylcholine (LPC), and LPC-O classes. Also, elevated levels of phosphatidylethanolcholines (PC), phosphatidylglycerols (PG), phosphatidylethanolserines (PS), and some phosphatidylethanolamines (PE) were found. Sphingomyelins (SM) were among the significantly affected sphingolipids (SL), exhibiting increased levels in similar patterns as GPLs (Fig. 3B, Table S4). Most of these lipids show an increasing trend throughout rats’ lifespan starting in 6-months with a gradual increase in 8 - 10-month-old animals and older (Fig. 3C). Lipid ontology (LION) enrichment analysis was used to characterize the lipidome changes from the biochemical and biophysical perspective (Fig. 3D). These results confirmed the significant involvement of GPLs, especially PCs and their lysoforms in tau-induced neurodegeneration. Membrane lipids with a positive charge headgroup accumulate in brain tissue, which may lead to alterations in membrane fluidity and affect endosomal/exosomal transport. High lateral diffusion, low bilayer thickness, and transition temperature point to elevated membrane fluidity [24].

**Fig.3.**
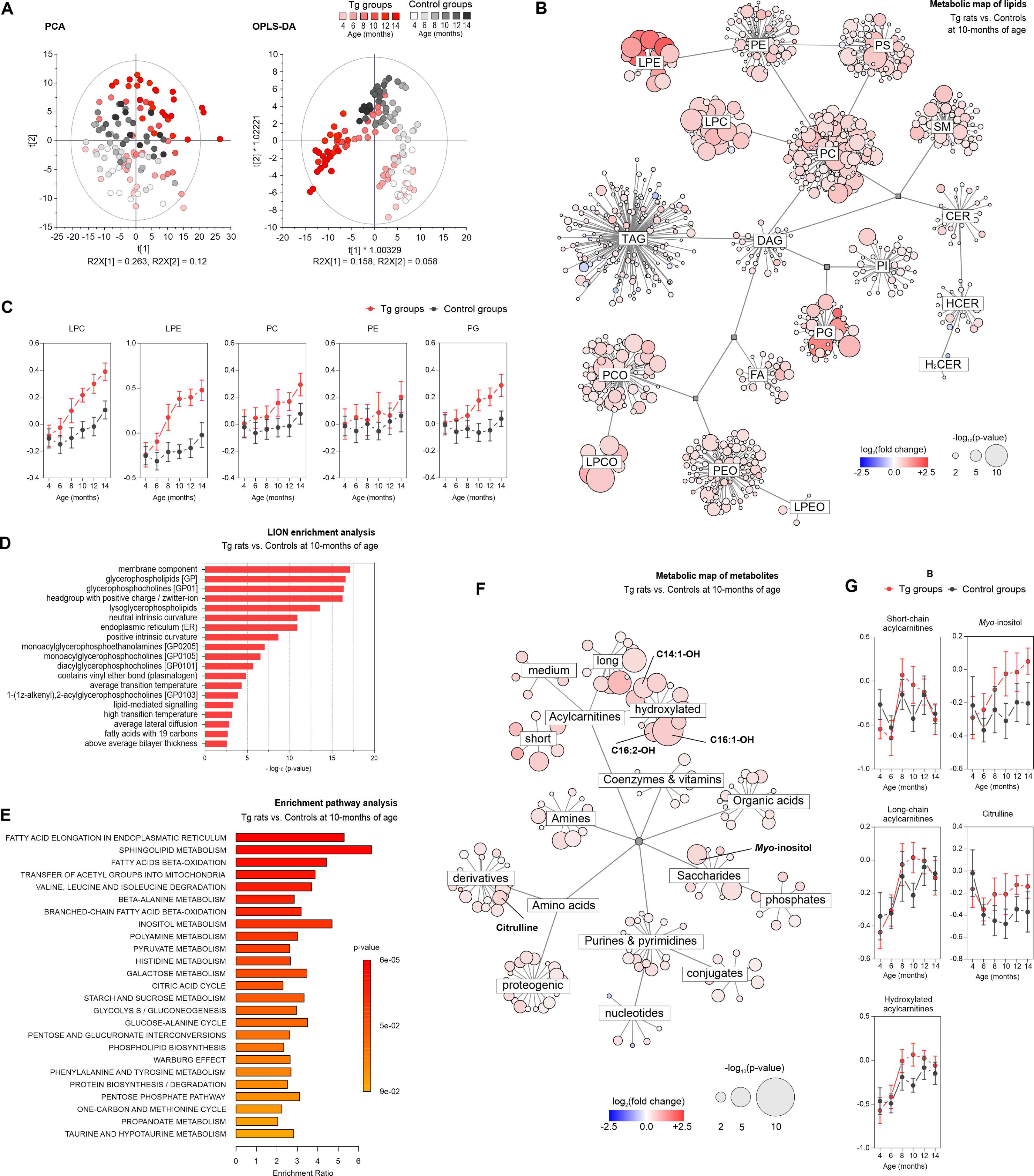
Metabolic patterns of brain tissue affected by tau pathology. Univariate and multivariate analyses of metabolic patterns found in the medulla oblongata of 10-month-old SHR-24 Tg rats compared with age-matched controls (SHR). **A** Principal component analysis (PCA) and orthogonal partial least square discriminant analysis (OPLS-DA) **B** Metabolic map showing results from significance testing for lipids – student’s t-test and fold change. The size of each bubble represents the statistical significance (p-value), and color and shade indicate the level of fold change between SHR-24 Tg and control groups. **C** Age-related changes of the most distinctive lipid classes (data are presented as median intensities with 95% confidence intervals). **E** Enrichment pathway analysis of metabolomics data. The biochemical pathways were assessed using statistical significance (p-value) and enrichment ratio. **F** Metabolic map showing results from significance testing for metabolites– student’s t-test and fold change. **G** Age-related changes of short-chain, long-chain, and hydroxylated acylcarnitines, myoinositol, and citrulline (data are presented as median intensities with 95% confidence intervals). . Explanation of lipid class abbreviations: LPE – lysophosphatidylethanolamines, PE – phosphatidylethanolamines, PS – phosphatidylethanolserines, PC – phosphatidylcholine, LPC – lysophosphatidylcholine, SM – sphingomyelins, CER – ceramides, HCER – hexosyl-ceramides, H_2_CER – dihexosyl-ceramides, DAG – diacylglycerols, TAG – triacylglycerols, PI – phosphatidylinositols, PG – phosphatidylglycerols, CE – cholesteryl esters, FA – fatty acids, PCO – plasmenyl phosphatidylcholines, LPCO – plasmenyl lysophosphatidylcholines, PEO – plasmenyl phosphatidylethanolamines, LPEO – plasmenyl lysophosphatidylethanolamines.

In contrast to gradual additive lipidosis seen in brain tissue lipidome, we observed subtle but very dynamic changes in the metabolome, mainly attributed to fatty acid elongation, and oxidation, transfer of acetyl groups into mitochondria, and sphingolipid metabolism (Fig. 3E, F, Table S5). Fatty acid (FA) and acylcarnitine (AC) metabolism were strongly affected in tau pathology. Aberrant patterns were observed in non-hydroxylated short- and long-chain, and hydroxylated acylcarnitines (Fig. 3G). While hydroxylated ACs followed the gradual increase in both groups, the levels in SHR-24 Tg animals preceded controls at 8 – 12 months of age. Long- and short-chain (including free carnitine) AC levels in the control group did not change during normal aging but significantly increased in 8-10-month-old Tg animals. We hypothesize a close association with the transition of tau into high-molecular tau aggregates which occurs around the 8^th^ month of age in Tg rats. A few hydroxylated ACs stand out, namely C14:1-OH, C16:1-OH, and C16:2-OH, that gradually increased with the disease progression in an additive manner. Also, elevated levels of citrulline, and *myo-*inositol, a known marker of glial activation [26–28] were found in the medulla oblongata of Tg rats compared to controls at 10-14 months of age (Fig. 3F, G, Table S5).

In pons, hyperphosphorylation and accumulation of aggregated forms of tau developed to a lesser extent and with a delay after medulla oblongata. Also, high-molecular tau aggregates (90 kDa) were mostly absent in pons, only appearing around 12 months of age. Lipid metabolism shows signs of lipidosis. Similarly, as in medulla oblongata, the most significant changes were observed in the levels of several lipids belonging to the following GPL classes: PC, PE and their lysoforms, PG and PS (Fig. S1B-C, and Table S4). Interestingly, the significant increase of these lipids was seen together with the appearance of high-molecular aggregates of tau protein at 12 months of age. Lipid ontology enrichment analysis in pons confirmed the importance of GPLs, especially lysoPLs in tau pathology-driven neurodegeneration (Fig. S1D). Similarly to medulla oblongata, we found very subtle changes in the pons metabolome of 10-month-old Tg rats (Fig S1E-F).

We concluded that pathological tau protein induced early lipid changes starting in 6-month-old Tg rats and gradually increased in 8 - 10-month-old animals and older. Our data showed a strong connection between the transition of tau into a sarkosyl insoluble fraction and changes in brain metabolism. The amount of insoluble tau protein corresponded to the extent of changes in lipidome and metabolome seen in the brain tissue of Tg rats.

### Cerebrospinal fluid, but not circulating blood, tracks the aberrant brain metabolism

The products of brain metabolism are secreted directly into the CSF and therefore, it is the main biological compartment for monitoring neurological processes. To investigate how tau-induced neurodegeneration is reflected in its surrounding biofluid, we performed metabolome and lipidome analysis of the CSF. PCA revealed substantial differences in metabolic profiles between Tg rats and controls. Groups according to age and neurofibrillary pathology were gradually separated. A clear separation appeared in 8-month-old animals and progressed with groups of 10, 12, and 14-month-old rats. The OPLS-DA analysis confirmed the PCA results with a clearer representation of the trends. (Fig. 4A).

**Fig.4.**
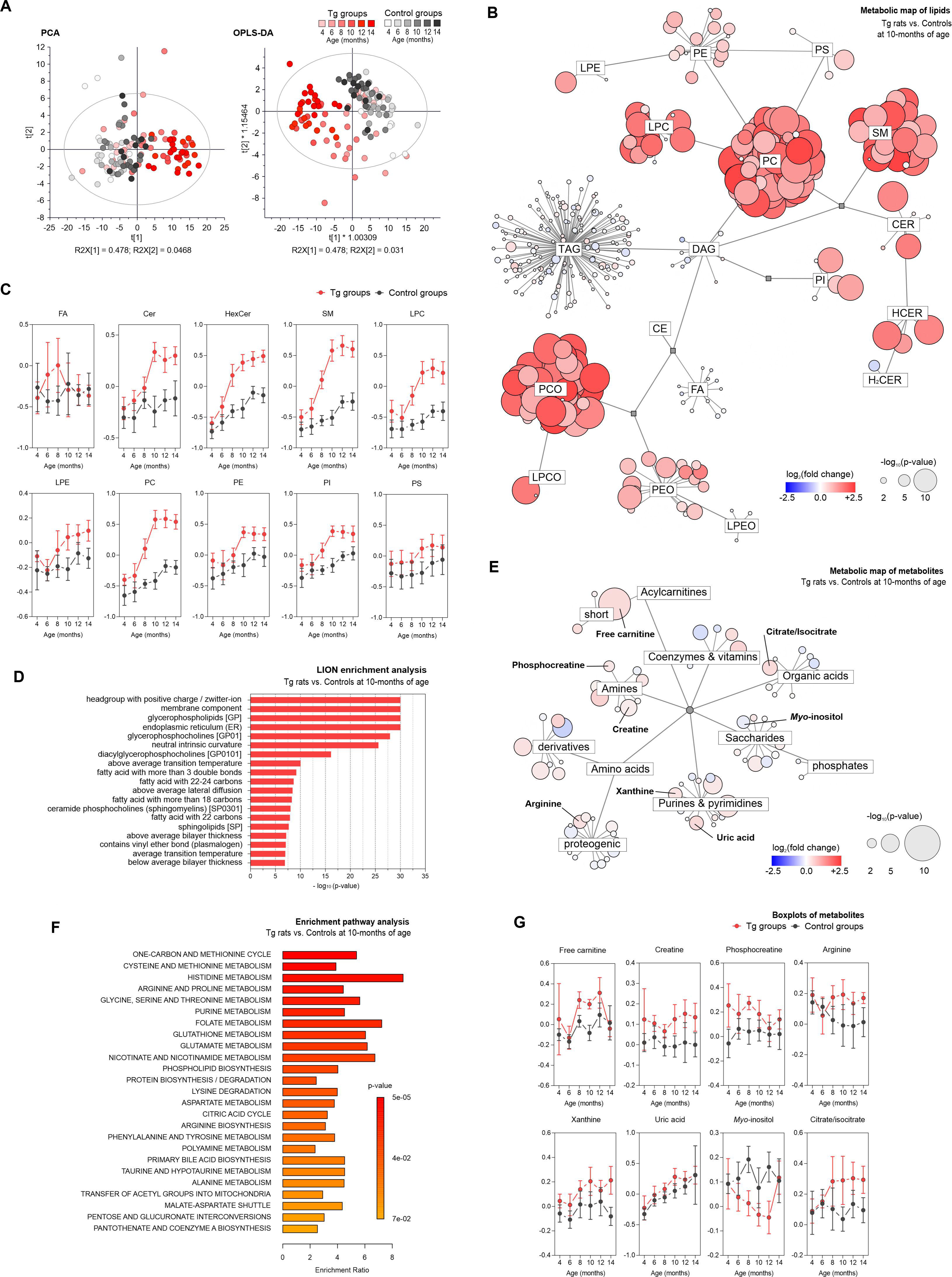
Global effects of brain aberrant metabolism on the composition of CSF. Univariate and multivariate analyses of metabolic patterns found in cerebrospinal fluid of 10-month-old SHR-24 Tg rats compared with age-matched controls. **A** Principal component analysis (PCA), and orthogonal partial least square discriminant analysis (OPLS-DA). **B** Metabolic map showing results from significance testing for lipids – student’s t-test (p-value and fold change). **C** Age-related changes of the most distinctive lipid classes (data are presented as median intensities with 95% confidence intervals). **D** Lipid ontology enrichment analysis. **E** Metabolic map showing results from significance testing for metabolites – student’s t-test and fold change. **F** Enrichment pathway analysis of metabolomics data. The biochemical pathways were evaluated using p-value and enrichment ratio. **G** Age-related changes of metabolites belong to the pathways: purine catabolism (xanthine, uric acid) and analytes of creatine/arginine metabolism (creatine, phosphocreatine, arginine), free carnitine, myo-inositol and citrate+isocitrate (data are presented as median intensities with 95% confidence intervals). Explanation of lipid class abbreviations: LPE – lysophosphatidylethanolamines, PE – phosphatidylethanolamines, PS – phosphatidylethanolserines, PC – phosphatidylcholine, LPC – lysophosphatidylcholine, SM – sphingomyelins, CER – ceramides, HCER – hexosyl-ceramides, H2CER – dihexosyl-ceramides, DAG – diacylglycerols, TAG – triacylglycerols, PI – phosphatidylinositols, PG – phosphatidylglycerols, CE – cholesteryl esters, FA – fatty acids, PCO – plasmenyl phosphatidylcholines, LPCO – plasmenyl lysophosphatidylcholines, PEO – plasmenyl phosphatidylethanolamines, LPEO – plasmenyl lysophosphatidylethanolamines.

Similarly, as in brain tissue, metabolic profiles in CSF were dominated by disturbed lipid metabolism. A significant number of aberrant lipids detected in brain tissue were reflected in CSF however, this was not an overall effect. A distinct pattern was observed in CSF, i.e. sphingolipids as elevated sphingomyelins and several ceramide (Cer) and hexosylceramide (HCer) species (Table S4). In the lipid profile, we have observed a significant increase in levels of GPL and SL, both categories similarly contributing to the CSF changes. Further univariate analysis revealed significant alterations in the levels of PC, PE, and their respective lysolipids. Similarly, as in brain tissue, CSF SM followed an increasing trend. A notably higher number of SMs were affected in CSF compared to brain tissue, and the differences between Tg and controls were more profound (Fig. 4B, C). In this study, we have also observed elevated levels of two ceramides (Cer 40:2 (d18:1/22:1), Cer 42:2 (d18:1/24:1)) and three hexosylceramides (HexCer 40:2 (d18:1/22:1), HexCer 42:2 (d18:1/24:1), HexCer 42:3 (d18:2/24:1)) species (Table S4). The dominant presence of membrane GPLs was suggested by lipid ontology (LION) enrichment analysis. Similarly, as in brain tissue, lipids with headgroups with positive charge were dominant in the CSF of Tg rats. The CSF lipidome was especially enriched in polyunsaturated fatty acids with 18 - 24 carbons and lipids containing ether bonds (Fig. 4D). Several metabolites involved in energy metabolism showed aberrant profiles in the CSF of Tg rats compared with controls (Fig. 4E). Elevated levels of free carnitine (between 8 – 10 months of age), creatine (from 6 – 14 months) and phosphocreatine (4^th^ and 8^th^ month) were found in the CSF. Interestingly, phosphocreatine was upregulated already in the youngest group of Tg rats, and the levels decreased with progressing neurodegeneration (Fig. 4G, Table S4).

The brain-specific isoenzyme creatine kinase, which ensures the conversion of creatine to its phosphorylated form, is highly susceptible to oxidative damage [29]. This could result in ongoing compensatory mechanisms or depletion of the antioxidant system. Elevated levels of xanthine, and uric acid may be another indicator of ongoing oxidative damage. Upregulated purine degradation pathway could be associated with increased production of radical oxygen species (ROS) [30], and development of neuroinflammation [31] (Fig. 4E, F, Table S4). Similarly to citrulline in the brain of Tg rats, elevated levels of arginine were found in the CSF. As opposed to the elevated levels in brain tissue, we have observed a decrease in the CSF *myo-*inositol in 8-12-months old Tg rats. Also, an increase in citrate+isocitrate was observed in 8-14-month-old Tg rats. These metabolites are associated with chronic neuroinflammation and the activation of glial cells [32,33] (Table S4).

Neurodegeneration often disrupts brain barriers and the presence of CNS molecules in the periphery [34]. Therefore, plasma samples were analyzed in this study to screen for possible peripheral markers of neurofibrillary pathology. The PCA model showed a partial separation of age-matched groups at the final stages of disease progression (10-14-month-old) (Fig. S2A). Surprisingly, CNS lipidosis was not reflected in plasma with only minor changes in several GPLs (Fig. S2B, Table S4). Plasma FAs were decreased during the early stages of disease progression up to the 10^th^ month of age. Also, reduced levels of several long-chain and hydroxylated acylcarnitines were found in plasma. However, this observation was limited to a 10-month-old group. Plasma-free carnitine was elevated in 8-10-month-old Tg rats. Several metabolites involved in arginine metabolism exert pathological patterns. Elevated creatine, decreased creatinine, and guanidinoacetate were detected in Tg rat plasma across most of the age-matched groups (Fig. S2C-E, Table S5).

### Pathological truncated tau secreted from neuron-like cells induced lipid changes in brain resident immune cells

Next, we aimed to study whether the extracellular pathological tau could induce lipid changes in glial cells. We used the multi-component cellular model system based on the co-cultivation of SH-SY5Y neuroblastoma cells inducible expressing pathological truncated tau and primary glial cells (microglia, astrocytes). The expression of truncated tau protein was induced by the removal of doxycycline from the cultivation medium.

At first, we analyzed the expression of endogenous and truncated tau in SH-SY5Y neuron-like cells (Fig. 5A, B). We quantified overexpression of total tau as well as tau phosphorylation on several AD-relevant epitopes (pSer262, pThr212, and pThr181) after 24, 48, 72, and 96 hours of induction of tau expression. We showed that in time the expression of truncated tau increased (Fig. 5D). On the other hand, the expression of endogenous tau was not significantly changed (Fig. 5C). We also showed that transgenic tau was more susceptible to phosphorylation compared to endogenous tau (Fig. 5B, D). The increased ability of truncated tau protein to be phosphorylated could be a result of different conformations compared to endogenous tau.

**Fig. 5.**
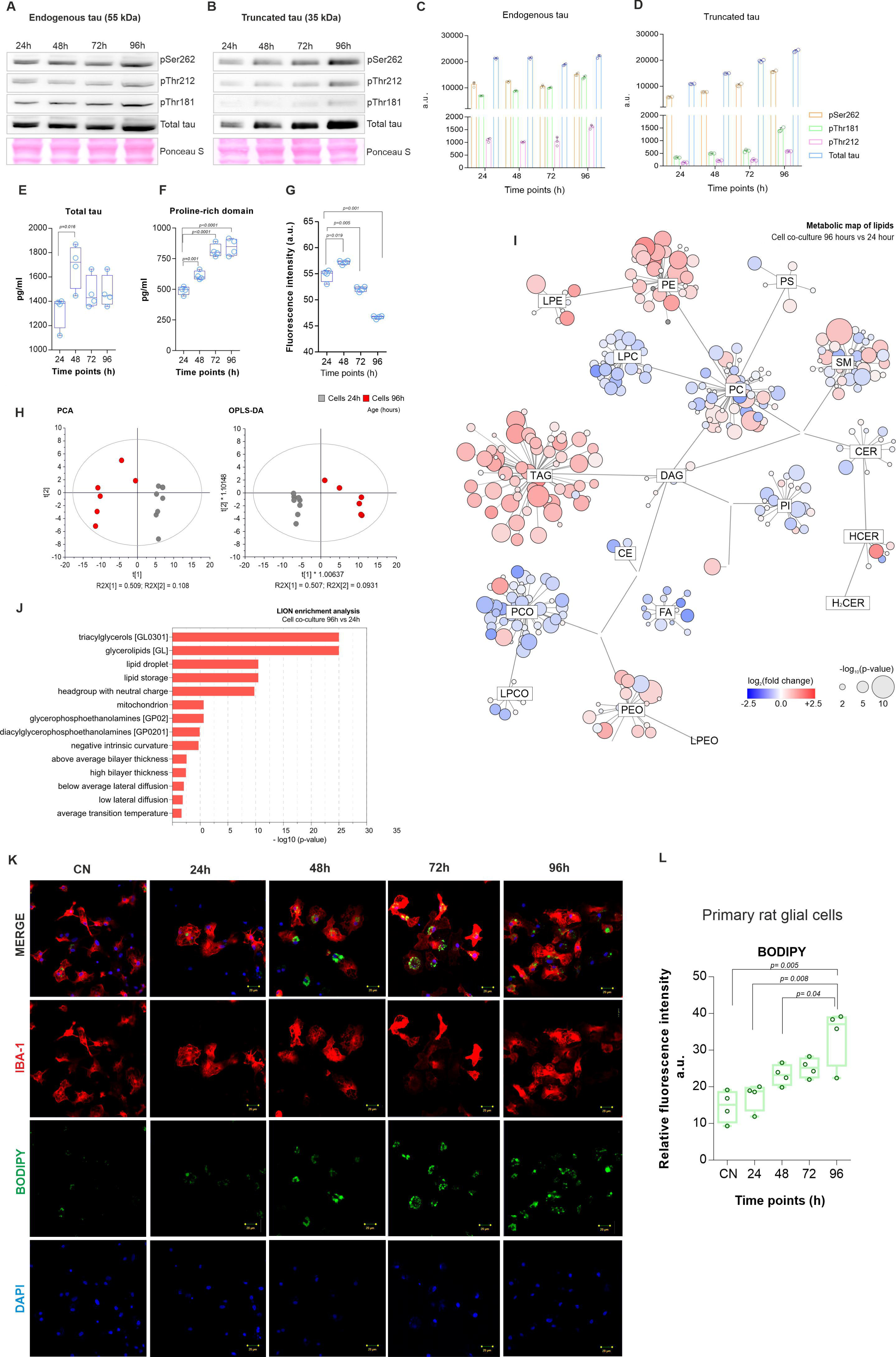
Pathological truncated tau protein induced lipid changes and LD formation in brain resident immune cells. **A-B** Expression of endogenous and truncated tau protein by human SH-SY5Y neuroblastoma cells. **C-D** Quantification showed that the expression of endogenous tau was not changed in time, however, truncated tau increased (mean±SD)**. E-F,** Analysis of cultivation media demonstrated higher secretion of the proline-rich domain of tau compared to total tau (boxplots with min to max, student’s t-test with p-value). **G** Cytotoxicity measured by AlamarBlue assay and displayed as boxplots (student’s t-test with p-value) **H** Principal component analysis (PCA), and orthogonal partial least square discriminant analysis (OPLS-DA) **I** Metabolic map showing results from significance testing for lipids – student’s t-test (p-value and fold change). **J** Lipid ontology enrichment analysis. **K-L** Primary rat microglia were co-cultivated with neuroblastoma cells expressing truncated tau. Representative images of DAPI (blue), BODIPY+ (green), and IBA-1 (red) staining in microglia and quantification of BODIPY+ relative fluorescence intensity (24 hours: 17.35 ± 1.84; 48 hours: 23.21 ± 1.4, 72 hours: 25.16 ± 1.33, 33.93 ± 3.9, n= 5; mean ± SD; student’s t-test with p-value) . Scale bar 20 µm.

Using ELISA, we quantified the amount of soluble monomeric tau released from truncated tau-expressing cells. Tau release has been detected in the cultivation media after 24, 48, 72 and 96 hours. We quantified the total tau and proline-rich domain of tau. We showed that after 48 hours the amount of total tau increased and then slightly decreased (Fig. 5E; 24 hours: 1322 ± 67.78 pg/ml; 48 hours: 1690 ± 89.48, 72 hours: 1471 ± 67.93, 96 hours: 1479 ± 65.67, n= 5). However, the amount of proline-rich domain of tau gradually increased in time (Fig. 5F; 24 hours: 491.2±16.5 pg/ml; 48 hours: 611.3±18, 72 hours: 815.2±26.4, 96 hours: 845.2±34.5, n= 5). To analyze if the overexpression of truncated tau affected cell viability, we performed an alamarBlue assay and detected a significant decrease in cell metabolic activity and proliferation after 72 hours compared to 24 hours (Fig. 5G; 24 hours: 53.54 ± 2.28; 72 hours: 52.08 ± 0.47, p=0.019, n= 4).

Lipidomic analysis was employed to evaluate how the lipid composition of the entire co-culture is influenced by the presence of tau released from neuron-like cells. Significant impacts were observed across almost the entire spectrum of measured lipids. There was a consistent increase in TAG, PE, and LPE classes, as well as a decrease in FA, CE, LPC, and LPCO classes. Additionally, there was an increase in SMs with shorter fatty acyl chains (up to 34 carbons) and a change in the regulation of PCs towards lower saturation levels (Fig 5I, J, Fig. S3). We also showed that tau protein secreted from neuroblastoma cells induced the formation of LDs in primary rat microglia. The formation of LDs was analyzed by immunocytochemical staining, and we found that the number of LDs accumulating cells significantly increased in time (Fig. 5K, L; 24 hours: 17.35 ± 1.84; 48 hours: 23.21 ± 1.4, 72 hours: 25.16 ± 1.33, 96 hours: 33.93 ± 3.9, n= 5). Based on the data, we propose that an increase in extracellular tau, particularly in the proline-rich domain in the media induces the formation of LDs in rat microglia.

To determine whether pathological tau protein triggers the formation of LDs, we also treated BV2 mouse-derived microglia with truncated tau (aa 151-391/3R). The formation of LDs was analyzed by immunocytochemical staining and flow cytometry (Fig. 6A - D). The results from immunocytochemistry showed a significant increase in the number of BODIPY^+^ cells (Fig. 6B, CN: 23.91 ± 5.62; Tau: 34.41 ± 8.60; p= 0.0013; n= 7) after tau treatment compares to control cells. Similarly, flow cytometry detected an increase in BODIPY mean fluorescence (Fig. 6D, CN: 1707 ± 227; Tau: 2325 ± 409; p= 0.0234; n= 4) compared to control conditions. Interestingly we showed that the tau protein changed the membrane fluidity of BV2 cells (Fig.6E, F). Pyrenedecanoic acid (PDA) was utilized as a reporter for cell membrane fluidity, due to its capability to undergo spectral shifting upon an increase in membrane fluidity. Changes in cell membrane lipid order between the gel/liquid-ordered phase (fluorescent at 430∼470 nm) and the liquid phase (fluorescent at 480∼550 nm), were determined by confocal microscopy. A shift to the liquid phase, indicating decreased membrane lipid order, was observed in BV2 cells treated with truncated tau protein. The membrane lipid order has not changed in control microglia cells (Fig. 6G). BODIPY-stained LDs were found to bind to mitochondria, suggesting that mitochondria regulate the processing of LD lipids (Fig. 6H). We investigated if the formation of LDs affected ATP levels. Luciferase luminescence ATP detection assay was performed, and we found decreased ATP activity in LD accumulating mitochondria (Fig. 6I, CN: 60346 ± 1310; Tau: 52267 ± 1032; p= 0.0029; n= 5).

**Fig. 6:**
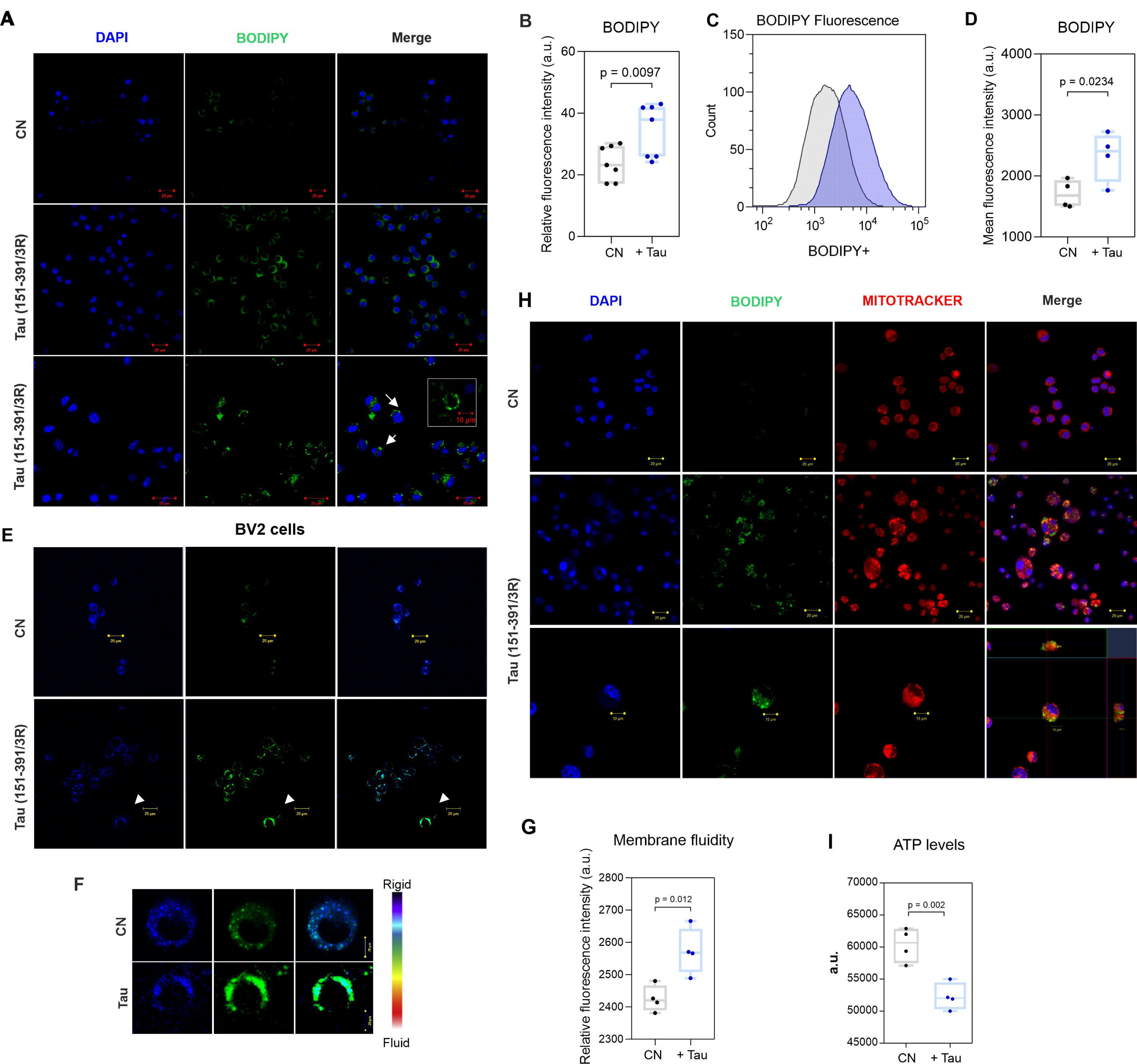
Tau protein-induced formation of lipid droplets and changes in membrane fluidity. **A** BV2 mouse-derived microglia were treated with Tau (151-391/3R) protein (500 nM for 24 hours). Representative images of DAPI (blue) and BODIPY+ (green) staining in BV2 cells. **B** Quantification of BODIPY^+^ relative fluorescence intensity (CN: 23.91 ± 5.62; Tau: 34.41 ± 8.60; p= 0.0013; n= 7; mean ± SD, student’s t-test with p-value). Scale bar 20 µm. **C** Representative flow cytometry histograms. **D** Quantification of BODIPY^+^ mean fluorescence intensity in BV2 cells (CN: 1707 ± 227; Tau: 2325 ± 409; p= 0.0234; n= 4; mean ± SD, student’s t-test with p-value). **E** Effect of Tau (151-391/3R) protein treatment on BV2 cells membrane fluidity. After incubation, BV2 cells were incubated with lipophilic pyrene probe PDA/Pluronic F127 solution (25 °C, 1 hour) which undergoes dimerization after the interaction and exhibits changes in its fluorescent properties. Changes in cell membrane lipid order, between the monomer gel/liquid-ordered phase (fluorescent at 430∼470 nm) and the excimer liquid phase (fluorescent at 480∼550 nm) were measured by confocal microscopy. **F** Representative images of changed membrane fluidity. **G** Quantification of relative fluorescence intensity of monomer vs. excimer ratio (n=4, mean ± SD; student’s t-test with p-value). **H** Representative images of DAPI (blue) and BODIPY+ (green), and MitoTracker (red) staining in BV2. Scale bar 20 µm. **I** ATP levels were decreased in LDs accumulating BV2 cells (CN: 60346 ± 1310; Tau: 52267 ± 1032; p= 0.0029, n= 5, mean ± SD; student’s t-test with p-value).

To confirm *in vitro* findings, we investigated LD numbers in microglia in Tg animals. Immunohistochemical analysis of rat brain tissue (Fig. 7A-B) showed a significant increase in LD-containing microglia (BODIPY^+^, IBA-1^+^) in the brainstem of Tg rats compared with control animals (Fig. 7B, CN: 1.2 ± 0.62; Tg: 3.5 ± 0.079; p= 0.0214; n= 10).

**Fig. 7:**
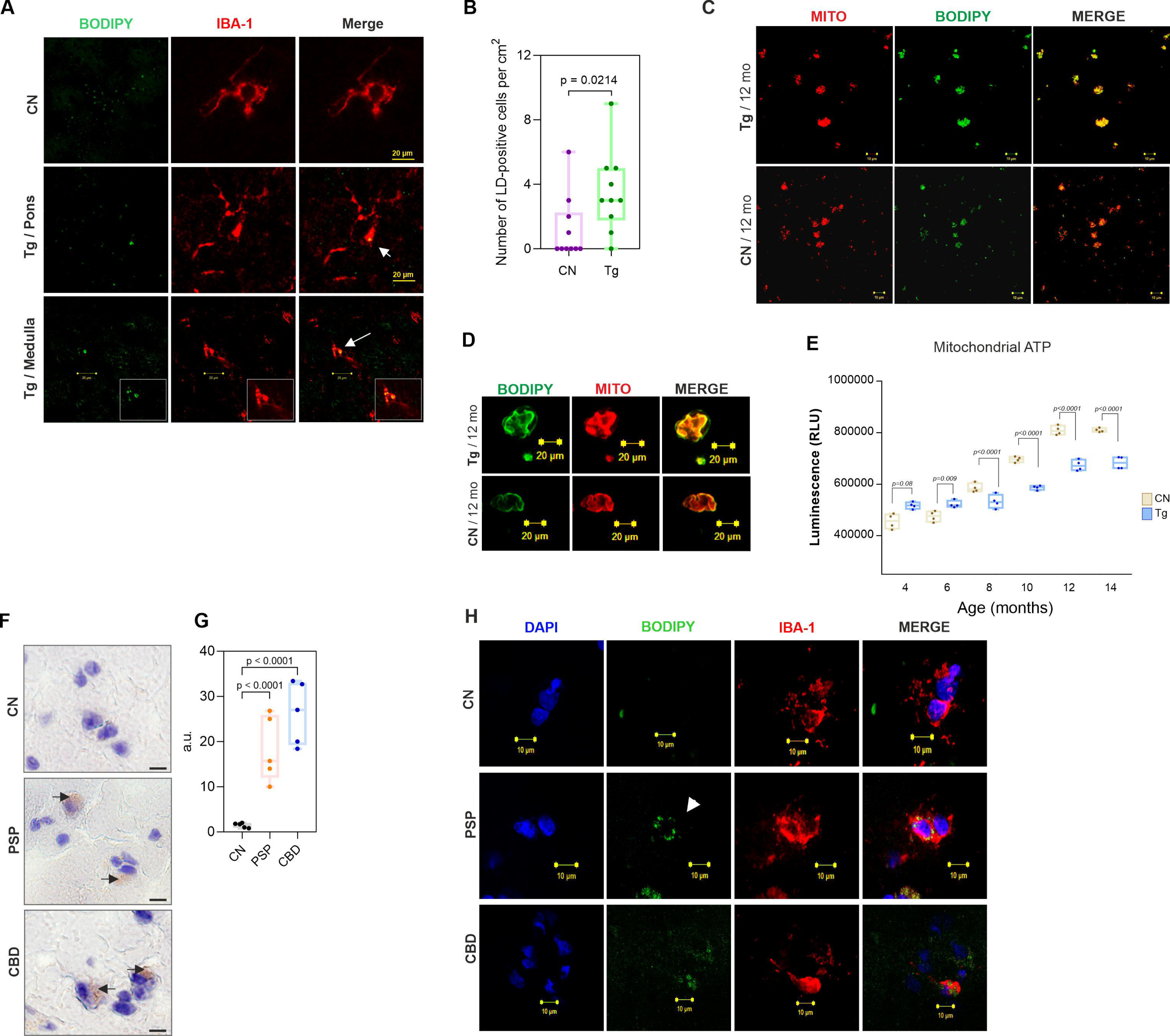
Lipid droplet accumulation in microglia of Tg rats and human tauopathies. **A** Representative confocal images. LDs accumulate in microglia of SHR-24 Tg rats. BODIPY^+^(green) and IBA1^+^(red) in the brainstem (medulla oblongata, pons) of 12-month-old SHR-24 Tg rats. Scale bar 20 µm. . **B** Quantification of BODIPY^+^ cells (CN: 1.2 ± 1.9; TG: 3.5 ± 2.5; p= 0.0214; n= 10, mean ± SD; student’s t-test with p-value). **C-D** LDM isolated from brain tissue of control and Tg animals. Representative images of BODIPY^+^ (green), and Mitotracker (red) staining in LDM. **E** Luciferase luminescence ATP detection assay showed decreased ATP levels in the LDM sample of Tg rats compared to controls. Data are presented as boxplots (Min to Max), student’s t-test with p-value. **F** LD formation in human brain tissue of patients with PSP and CBD. Oil Red O staining neutral lipids images. **G** Quantification (CN: 1.46 ± 0.53; PSP: 18.30 ± 7.27; p < 0.0001 CBD: 26.30 ± 6.96; p< 0.0001; n= 5, student’s t-test with p-value). **H** Representative images of DAPI (blue), BODIPY^+^ (green), and IBA^+^ (red) staining in the human brain. Scale bar 10 µm.

Since LDs and mitochondria are organelles involved in cellular lipid metabolism and energy homeostasis we isolated lipid droplet-associated mitochondria (LDM) from the brain tissue of transgenic and control rats. Similarly to immunocytochemistry, LDs were associated mainly with mitochondria from Tg animals (Fig. 7C, D). Further, a luciferase luminescence ATP detection assay was performed, and we find decreased ATP levels in the LDM samples from Tg rats compared to controls. We have detected a significant decrease in ATP levels already at 8 months of age (Fig. 7E, p=0.016, n=4). We conclude that the accumulation of LDM negatively affects cell bioenergetics.

Moreover, we analyzed the formation of LDs in human brains from two different tauopathies. Oil red O, together with fluorescence BODIPY^+^, IBA-1^+^ staining was used to stain LDs in human brain tissue from PSP and CBD patients. We have observed a significant accumulation of LDs in human brain microglia (Fig. 7F-H). Together, these findings demonstrated that tau protein triggers LD formation in brain resident immune cells *in vitro* and *in vivo*.

### Tau pathology causes a switch from glucose metabolism to fatty acid oxidation

Lastly, we examined whether neurofibrillary pathology changes primary energy metabolism. First, we analyzed the expression of glucose transporter 3 (GLUT3) in the medulla oblongata and pons of Tg animals and age-matched controls (Fig. 8A-B). We determined the levels of GLUT3 by immunofluorescence analysis. As a neuronal marker, we used the neuron-specific protein NeuN. No changes were found in 4-month-old animals (Fig.8D, 4-month-old animals: CN: 32.07 ± 6.1, Tg: 35.01 ± 5.23). We observed the first changes in 6-month-old Tg animals. Quantification showed NeuN-positive neuronal cells with slightly decreased GLUT3 staining (Fig. 8B, 6-month-old animals: CN: 28.04 ± 2.4, Tg: 29.36 ± 2.1; white arrows). Further quantification showed that the decrease of GLUT3 was more significant in 8-, and 10-month-old animals (Fig.8C, D, 8-month-old animals: CN: 25.07 ± 1.43, Tg: 20.91 ± 0.54; p= 0.035; 10-month-old animals: CN: 28.37 ± 1.03, Tg: 22.36 ± 1.67; p= 0.022). This points out that the decrease in neuronal GLUT3 is most likely specific to neurofibrillary pathology rather than being caused by neuronal loss.

**Fig. 8:**
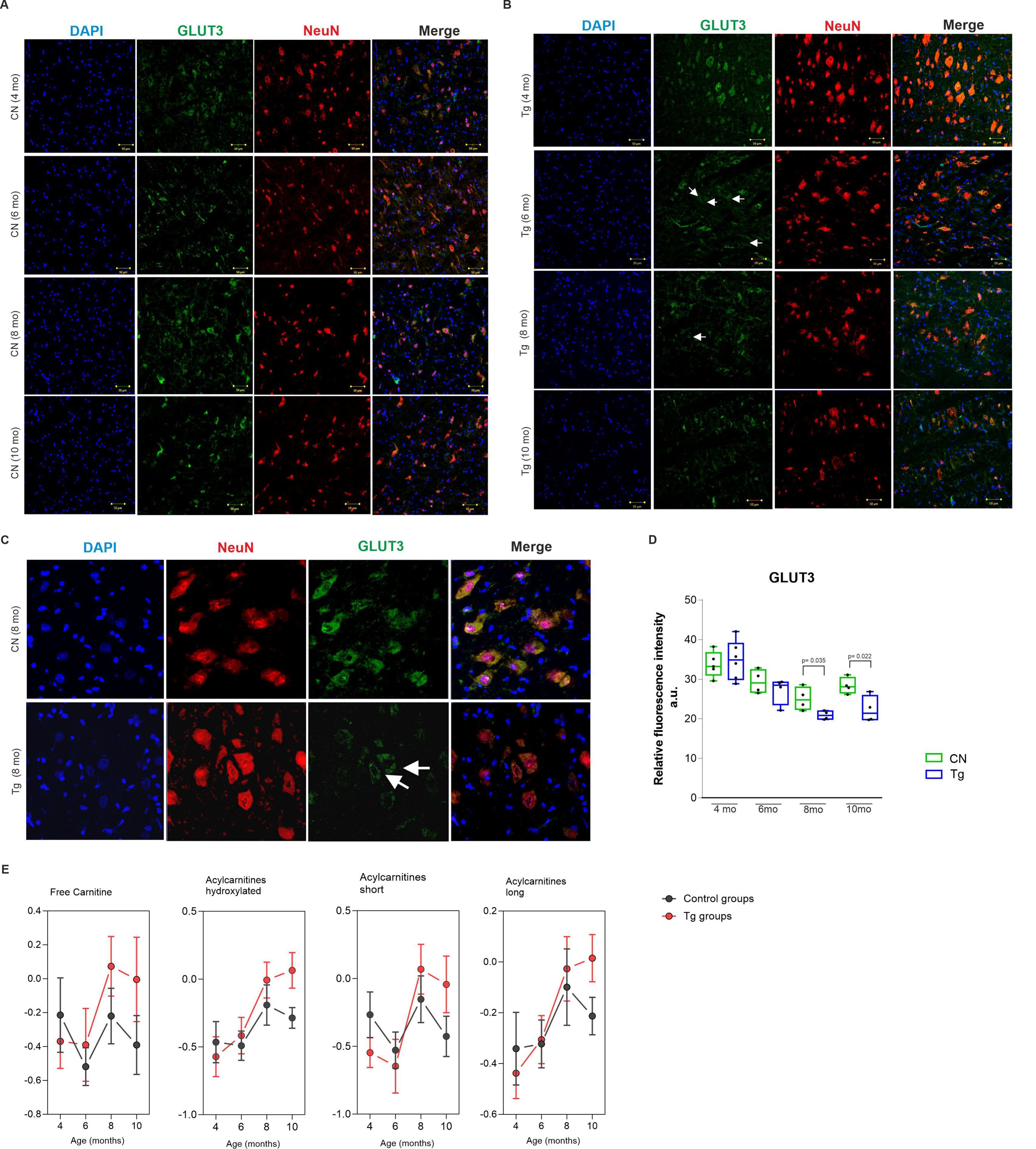
Tau pathology switches the energy metabolism to fatty acid oxidation. **A-B** Immunofluorescence staining of neuron-specific glucose transporter 3 (GLUT3). Representative images of GLUT3 (green), NeuN (red), and DAPI staining in brain tissue from control and transgenic animals. **C** Representative images of 8-month-old control and transgenic animals. Scale bar 20 µm. **D**. Quantification of GLUT3 mean fluorescence intensity in brain tissue. Quantification of relative fluorescent intensity showed a decrease of GLUT3 in transgenic animals (8-months old animals: CN: 25.07 ± 1.43, Tg: 20.91 ± 0.54; p= 0.035; 10-month-old animals: CN: 28.37 ± 1.03, Tg: 22.36 ± 1.67; p= 0.022; n= 5; mean ± SD, student’s t-test with p-value). **E** Changes of long-chain, hydroxylated, short-chain ACs and free carnitine in brain tissue of control and transgenic rats (data are presented as median intensities with 95% confidence intervals).

To study whether the decrease in GLUT3 levels induces the shift towards fatty acid metabolism we determined the production of short-chain, long-chain, hydroxylated ACs as well as free carnitine (Fig. 8E). Free carnitine as a neuroprotective molecule already increased in 6-month-old Tg rats compared to controls. Long-chain and hydroxylated ACs significantly increased in 8-month-old Tg animals. Interestingly, after 8 months we also observed a mild increase in short-chain ACs. It suggests that the decrease in GLUT3 may contribute to the high amount of long-chain and hydroxylated ACs in the brain tissue of Tg rats. Taken together, these results suggest that after GLUT3 decrease, an increase in ACs contents in the brain tissue of Tg rats is observed as a result of the energy metabolism switch from glucose to fatty acid oxidation.

## Discussion

The pathological aggregation of tau protein is a hallmark of neurodegenerative diseases collectively referred to as tauopathies. Growing evidence suggests the significant involvement of deregulated lipid metabolism in the pathogenesis of neurodegenerative diseases, including tauopathies. While the relationship between lipid metabolism and tauopathy is not as well understood as that between lipid metabolism and Aβ pathology, recent studies have indicated a potential link between lipid metabolism and abnormal tau aggregation [35]. In the present study, targeted lipidomic and metabolomic analysis was employed to reveal metabolic alterations induced by the cascade involving tau pathology.

Using a transgenic model with progressive age-dependent neurodegeneration and well established multi-component cell model system, we demonstrated that misfolding of tau is associated with marked metabolic changes. The presence of extracellular and aggregated tau was associated with increased production of lipids participating in protein fibrillization, membrane reorganization, and inflammation. Consequently, the membrane fluidity in glial cells was affected, leading to the accumulation of lipid particles in mitochondria, thus affecting the cell bioenergetics of brain cells.

Several independent studies characterized the neurolipidome of the AD human brain and identified changes in the abundance of a number of lipid species [28, 32]. In the current study, dysregulation of lipid metabolism was prominent in CSF and brain tissue (pons, medulla) of tau Tg rats compared to age-matched controls. Comprehensive lipidomics analysis established that LPC, LPE, PC, and PG were increased in brain tissue and CSF of Tg rats compared with controls. We were also interested in understanding the lipid dysregulation associated with disease progression and analyzed the transgenic and control animals in longitudinal samples (six-time points). Interestingly, significant changes have been found in the early steps of tau aggregation before the formation of sarkosyl-insoluble high-molecular-weight aggregates. The altered levels of several classes of GPL metabolism (LPC and LPE) were increased with time in Tg animals compared with controls. These observations suggest that lysophospholipid (lysoPL) species (LPC and LPE) are particularly associated with disease progression. GPLs normally are components of cellular or vesicle membranes. Membrane lipids act as essential signaling molecules and key modulators of signal transduction and vesicle trafficking [36,37]. LysoPLs are intermediate metabolites in well-regulated phospholipid biosynthetic pathways produced by phospholipase-mediated hydrolysis [38]. Aß aggregation has been shown to induce PLA2 up-regulation and activation [39], and increased levels of lysoPLs have been found in brain ischemia [40], traumatic brain injury [41], and amyloid transgenic mice [42]. LPE and LPC bind to proteins such as receptors, enzymes (kinases, phosphatases), and act as neurotrophic factors that promote neuronal differentiation. The decrements observed in our Tg rats could indicate a potential role in dysfunctional synaptic transmission early in the disease process. The phospholipid degradation to lysoforms could also play an important role in the early stages of tau accumulation. We found significantly increased amounts of LPC and LPE in 6-months old animals before the development of high-molecular-weight tau aggregates. This suggests that LPC and LPE classes could be crucial for the early steps of tau aggregation and can promote further protein fibrillization. Previously, LPC has been shown to directly enhance the formation of Aß(1-42) fibrils [43,44]. Except for protein fibrillization, the accumulation of lysoPLs in brain tissue elicits neurotoxic effects, through the release of proinflammatory molecules [45], astrocyte and microglial activation [46], and alterations in membrane organization/fluidity [47]. Our previous work showed that abnormal truncated tau induced inflammatory response in later stages of pathology [48], and therefore the gradual increase of LPC and LPE in brain tissue of Tg rats could be the result of progressive neuroinflammation.

During the early stages of tauopathy, we showed significant upregulation of pathological hyperphosphorylated forms of tau protein and increased levels of total tau in the CSF of Tg rats. We, therefore, anticipate that lipid deregulation during the early stages of tauopathy could be caused by increased secretion of extracellular tau. To better understand how extracellular tau modulates lipid metabolism, we used a multi-component *in vitro* cell model system. The model was developed by co-cultivation of human neuroblastoma cells overexpressing pathological truncated tau and primary rat glial cells (microglia, astrocytes). We showed that human neuroblastoma cells overexpressing truncated tau actively release monomeric total tau and proline-rich domain of tau into the media. Interestingly, levels of secreted proline-rich domain of tau significantly increased in time. A lipidomic study revealed increased production of lipids participating in protein aggregation a membrane reorganization as well as the formation of Soluble truncated tau protein released by neuron-like cells induced the formation of LDs in microglia. Our results showed that this process depended on the amount of secreted tau. Tau-mediated accumulation of LDs was also shown in the Tg rat model and on human brain tissue from patients with tauopathies. LDs were found adjacent or partially colocalized with mitochondria and this was associated with decreased ATP activity. The formation of LDs could be caused by the accumulation of lipidic by-products of inflammatory response that cannot be processed due to dysfunctional mitochondria. Little is known about the dynamics of LD composition, which depends on the cell type and its current metabolic state [49]. Previous reports showed that immune cells such as microglia accumulate large amounts of lipids in LDs [50,51]. LD-accumulating microglia are defective in phagocytosis and secrete high levels of proinflammatory cytokines. LDs have been previously proposed to serve as a lipid buffering system to prevent acylcarnitine accumulation and lipotoxicity in mitochondria [52]. <u>The</u> lipid dyshomeostasis within microglia is likely to alter neuron-glia interplay and the metabolic integrity of neurons, leading to accelerated neurodegeneration.

Previous evidence has demonstrated that full-length or truncated tau is secreted by neurons and actively released into extracellular space. Although the physiological function of extracellular tau remains unclear, it is suggested its role in trans-cellular propagation. Extracellular tau can spread through the brain tissue and be (1) internalized by healthy neurons affecting their function and (2) activate glial cells during neuroinflammation [53,54]. Tau secretion involves multiple pathways including vesicle-free direct translocation across the plasma membrane (PM). It was shown that during pathological and physiological conditions, the majority of tau is secreted in a vesicle-free form via translocation across PM. Tau filaments were found at the PM of AD brains. Membrane lipids such as PC and sphingolipids were associated with tau filaments isolated from AD patients. It was confirmed that MTBD of tau is involved in the binding of tau to lipid membrane and N-terminal fragment of tau containing proline-rich domain promotes tau aggregation. Based on the data from transgenic animals and *in vitro* model systems, we showed that the early lipid changes before the development of neurofibrillary pathology could be induced by overexpression and increased secretion of extracellular tau. Our data showed that in the early stage of tauopathy, extracellular tau caused dysregulation of lipid homeostasis, which could accelerate further neurodegenerative and inflammatory processes. The subsequent abrogation of lipid homeostasis could lead to tauopathy-like molecular phenotypes.

During the further development of neurofibrillary pathology, we uncovered the deregulation of SM metabolism. A significant increase in the levels of several SM, Cer, and also hexosyl-ceramide (HexCer) species was detected. Increased levels of SM in CSF of Tg rats could be caused by membrane breakdown, demyelination, and progressive decline of neuronal cells in the brain tissue during the development of neurofibrillary pathology. SM metabolism is tightly associated with the metabolism of ceramides (Cer) [55]. We observed elevated levels of two specific ceramides (Cer 40:2 (d18:1/22:1), Cer 42:2 (d18:1/24:1)) and three hexosylceramides (HexCer 40:2 (d18:1/22:1), HexCer 42:2 (d18:1/24:1), HexCer 42:3 (d18:2/24:1)) species. Several studies connect aberrant Cer production to Aß pathology [56,57]. Aß oligomers cause activation of SMase enzyme, leading to Cer accumulation and subsequent cell death [58]. In AD, levels of SM in CSF correlate with the levels of Aß and tau protein, and elevated levels of very-long-chain Cer species were detected in APOE [4 carriers [59].

With age and pathology progression also several phosphatidylglycerol (PG) species have been gradually elevated in the brainstem of Tg rats. PG is known mainly for its function as a pulmonary surfactant, however in mammalian cells, PG is synthetized in mitochondria and involved in processes such as RNA transcription, and activation of protein kinase C (PKC) [60]. In a 5XFAD mouse model of AD, increased activity of PKC was found in astrocytes adjacent to Aß plaques [61]. Interestingly, PKC requires phosphatidylserine (PS), localized on the cytoplasmic leaflet for its activation [62]. Alongside PG accumulation we also observed elevated levels of PS species in brain tissue, however to a lesser extent. This is in good agreement with studies on human AD brain tissues [63]. PKC can promote tau protein phosphorylation [64]. Therefore, pathological PKC activation by PG and PS lipids may be involved or further promote tau hyperphosphorylation and progressive formation of neurofibrillary tangles in Tg animals.

PCs and PEs, as the two most abundant membrane phospholipids across all types of cells, were found to be significantly increased in the brain tissue of 8 months old rats. These lipid species could convoy inflammatory processes associated with widespread neurofibrillary pathology. Both elevated and decreased levels of PCs were found in the APP mouse model for late-onset AD [65]. However, no significant alterations [66], or even decreased levels [67] were found in post-mortem AD brain tissue and human plasma of AD patients [68].

The findings from the metabolic study showed that significant changes occur in the early stages of diseases before the formation of sarkosyl-insoluble high-molecular-weight tau aggregates. We found that tau accumulation leads to disruption in metabolic pathways associated with mitochondria, such as fatty acid oxidation (FAO), elongation, tricarboxylic acid cycle (TCA), creatine metabolism, purine catabolism, and arginine metabolism. These results are consistent with the observation that soluble tau plays a role in mitochondria distribution and oxidative stress, and the long-term mitochondrial stress can trigger further tau dimerization [69,70].

We found changes in creatine and phosphocreatine in 4- and 6-month-old Tg rats compared to controls. Although we found no changes in brain creatine (Cr) nor phosphocreatine (PCr) in brain tissue, we observed their increased levels in the CSF. The Cr/PCr system involving brain-specific Cr kinase (CK) enzyme, functions as an energy buffer in the ATP-mediated cellular energetics [71]. Disturbances in Cr/PCr balance are indicative of impaired energy metabolism in mitochondria and oxidative stress during the early stages of pathology in Tg rats. Brain-specific CK activity has been shown to decrease in AD brain [29].

Our analysis also showed that the transition of tau protein from monomeric to high-molecular-weight oligomers reprograms the brain metabolism from glycolysis to FA uptake and beta-oxidation. Recent evidence shows that FAO plays a crucial role in generating energy in mitochondria [72,73]. However, brain energy metabolism is prevented from using FAs as a source of energy for their cytotoxicity and potential capacity to stimulate neurodegeneration [74,75]. We showed that expression of the neuron-specific glucose transporter GLUT3 was decreased in Tg rats, and this positively correlated with the amount of neurofibrillary pathology in brain tissue. In the case of impaired glucose uptake and metabolism, deregulated catabolism of FAs serves as a regulator of cell energy metabolism. Metabolism of FAs and glucose yield ACs, so their concentration is determined by the nutritional state. Short-chain ACs are produced from glucose, amino acids, and FA degradation, medium- and long-chain ACs are derived from FA metabolism. Long-chain ACs are synthesized in mitochondria; therefore, long-chain ACs are used as markers of mitochondrial FA oxidation. Our results showed increased levels of hydroxylated and non-hydroxylated long-chain ACs in the brain tissue of Tg rats. Their amount increased from 6 to 12-month-old Tg animals compared to controls. The increase coincided with the appearance of tau aggregates in brain tissue. Increased levels of hydroxylated and non-hydroxylated long-chain ACs were also caused by the aggregation of amyloid-β in brain tissues of APP/PS1 transgenic mice models [76]. On the other hand, the amount of short-chain ACs decreased in 4 and 6-month-old Tg rats. Interestingly, their levels increase from 8 to 10-month-old Tg animals. Our result also showed that the brain levels of ACs returned to control levels, except for C14:1-OH, and C16:1-OH, which remained elevated up to 14 months of age. This suggests that the brain affected by tauopathy can, to some extent, stabilize its metabolic status.

Except for the mentioned ACs, elevated levels of free carnitine were found in brain tissue, CSF, and plasma. However, this upregulation was limited to 8-10-month-old tau Tg animals, which may be associated with higher demand for lipid catabolism. Free carnitine is reported to have neuroprotective effects [77,78] and provides the transport of FAs into mitochondria by the carnitine shuttle system. The exceptions are short- and medium-chain FAs, which can diffuse across both mitochondrial membranes. These transport systems are involved in maintaining proper function and regulation of the elemental metabolic pathways such as the TCA cycle and FAO [79]. Lipid accumulation leads to an imbalance between these two crucial pathways and manifests as an increase in long-chain ACs [80,81], which is also demonstrated by our metabolomic data.

## Conclusion

In conclusion, our results showed that misfolding and aggregation of tau lead to metabolic changes that are different in different stages of tau pathology. Our study provides new evidence that support the contribution of pathologic tau proteins in individual lipidome pathways. The lipids play an important role in neuron-glia communication and can modulate further processes. During the early stages of tauopathy, lipid homeostasis is disturbed by increased secretion of extracellular tau, which could accelerate neurodegeneration. The first steps of tau protein aggregation are associated with metabolites and lipids playing an important role in energy metabolism, membrane integrity, inflammation, and cellular signaling. The results support a strong connection between the transition of tau into a sarkosyl insoluble fraction and changes in brain metabolism. Based on the data we believe that biologically active membrane lipids such as phospholipids and sphingolipids could represent new potential therapeutic targets in neurodegenerative diseases such as tauopathies.

## Supporting information

Olesova et al supplementary

## Acknowledgments

This work was supported by a Doctoral student grant (DSGC-2021-0098) from University Palacký Olomouc, MH CZ - DRO FNOl, 00098892 from University Hospital Olomouc, APVV-21-0321, APVV-22-0313, and VEGA 2/0129/21.

## Author contributions

The supervision and design of this study were carried out by A.Ko. and D.F. Sample collection was done by A.M., B.J., D.O. Sample preparation was done by R.B., D.O., D.D., and Š.K.

LC-MS analysis was performed by D.O., D.D., D.F., and Š.K. Western blot and other biological experiments were done by P.M., J.P, J.H., L.F., and E.S. Data treatment and statistical analysis were carried out by R.B., A.Kv., D.D. and D.F. Visualization and interpretation of the results were done by D.D., D.O., A.Kv., D.F. and R.B. The first version of the manuscript was writing by D.O. and D.D. This version was edited and reviewed by all co-authors.

## Ethical approval

All animal experiments were performed according to the institutional animal care guidelines and in accordance with international standards (Animal Research: Reporting of In Vivo Experiments guidelines) and approved by the State Veterinary and Food Administration of the Slovak Republic (Ro-933/19-221/3a) and by the Ethics Committee of the Institute of Neuroimmunology, Slovak Academy of Sciences. Human brain tissue samples were obtained from the Queen Square Brain Bank for Neurological disorders (London, UK). All human samples were obtained with informed patient consent and with approval from the local ethical committees.

## Data availability

The attachments presented in this study are included in the Supplementary material. Other questions can be directed to the corresponding authors.

Lipidomic and Metabolomic data have been submitted to MassIVE repository, registered under ID : MSV000091311 [doi:10.25345/C5D50G70W], and can be accessed using the download link: ftp://massive.ucsd.edu/MSV000091311/.

## Abbreviations

AC: Acylcarnitine
AD: Alzheimer’s disease
CNS: Central nervous system
CSF: Cerebrospinal fluid
FA: Fatty acid
GLUT3: Glucose transporter 3
LD: Lipid droplet
LDM: Lipid droplet-associated mitochondria
LPC: Lysophosphatidylcholine
LPE: Lysophosphatidylethanolamine
NFL: Neurofilament light chain
SM: Sphingomyelin
TGA: Triglycerides

