## Supplementary figures and images for "Changes in lipid metabolism track with the progression of neurofibrillary pathology in tauopathies"

### FigS1.jpg

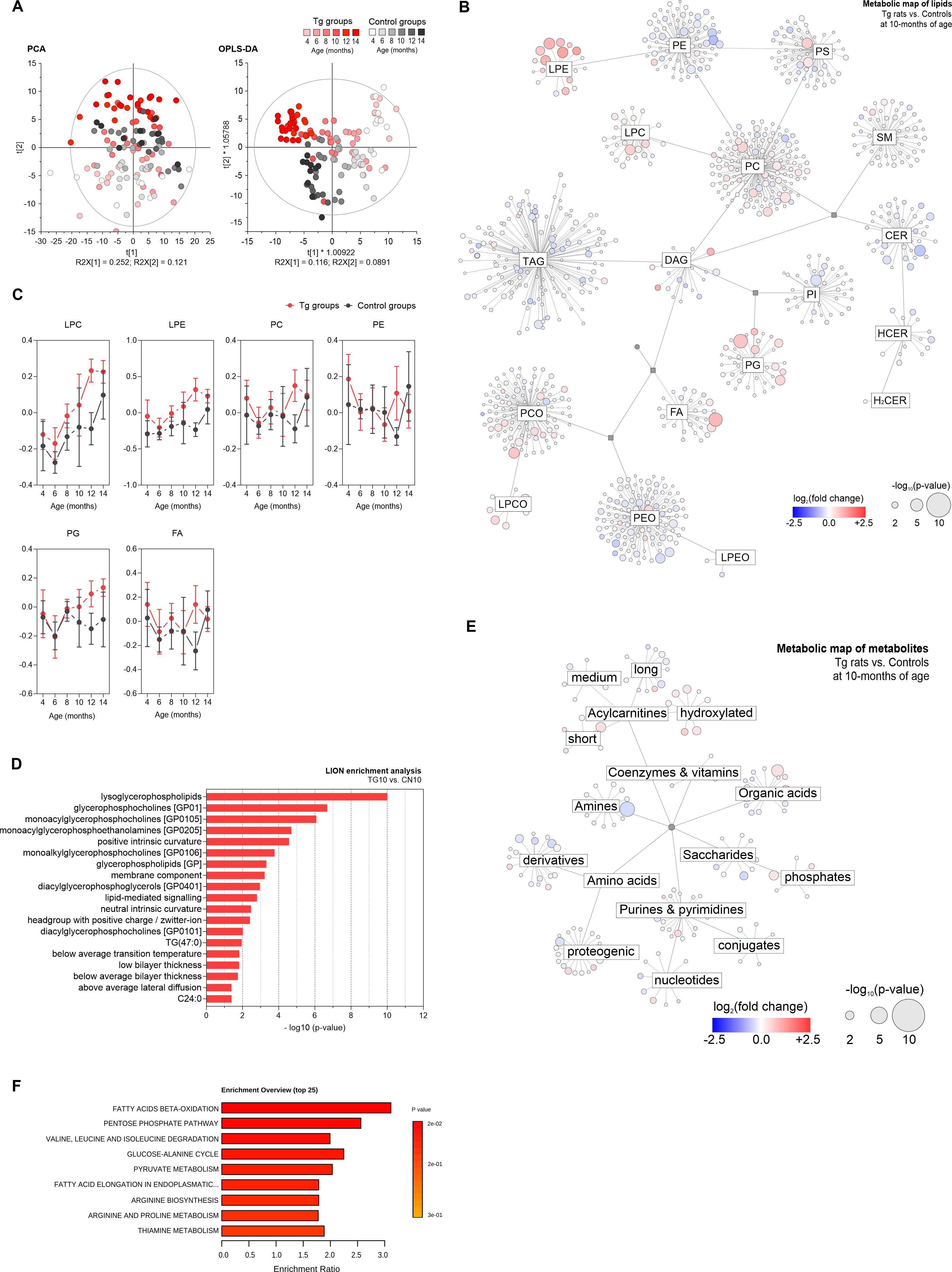

### FigS2.jpg

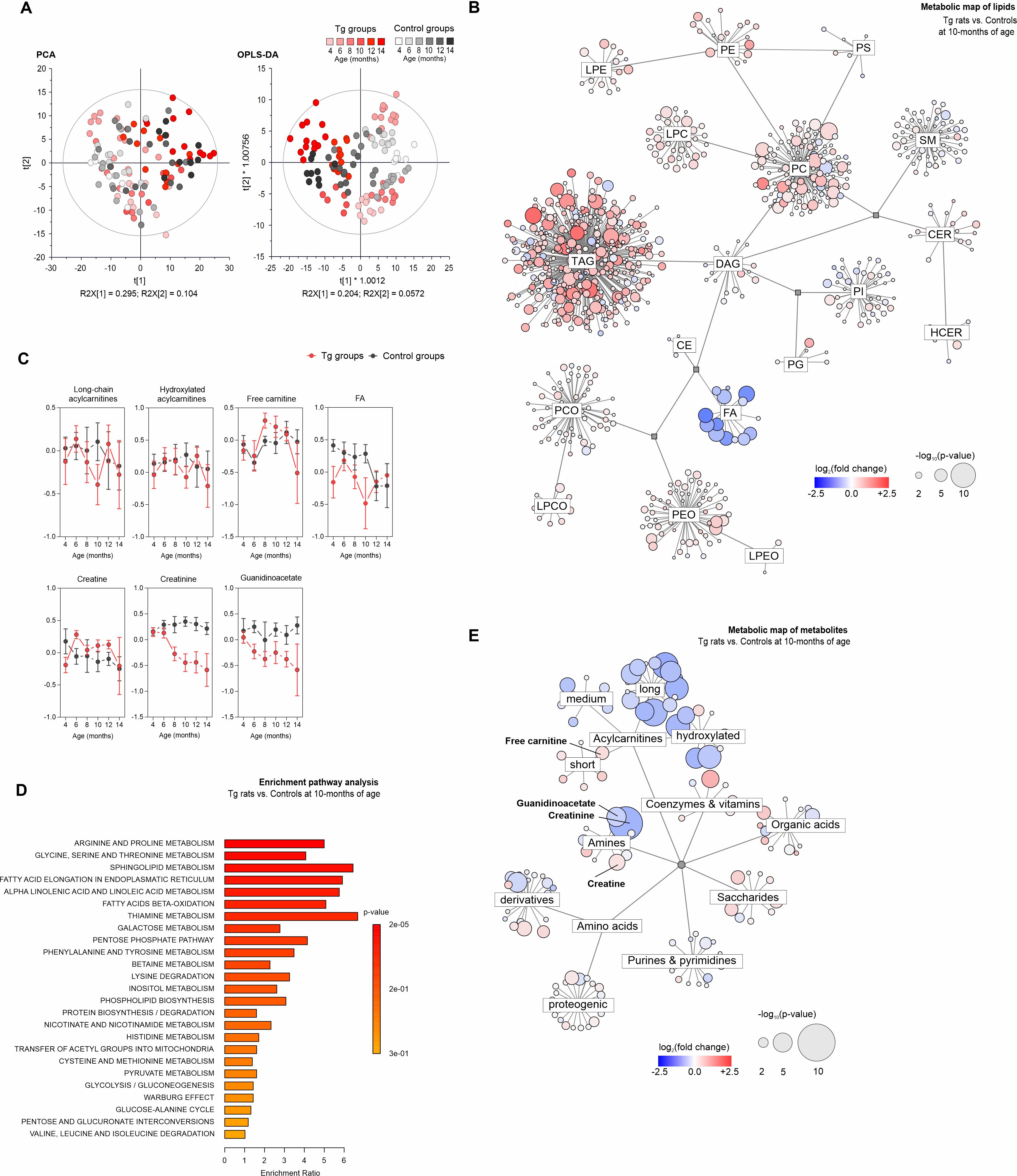

### FigS3.jpg

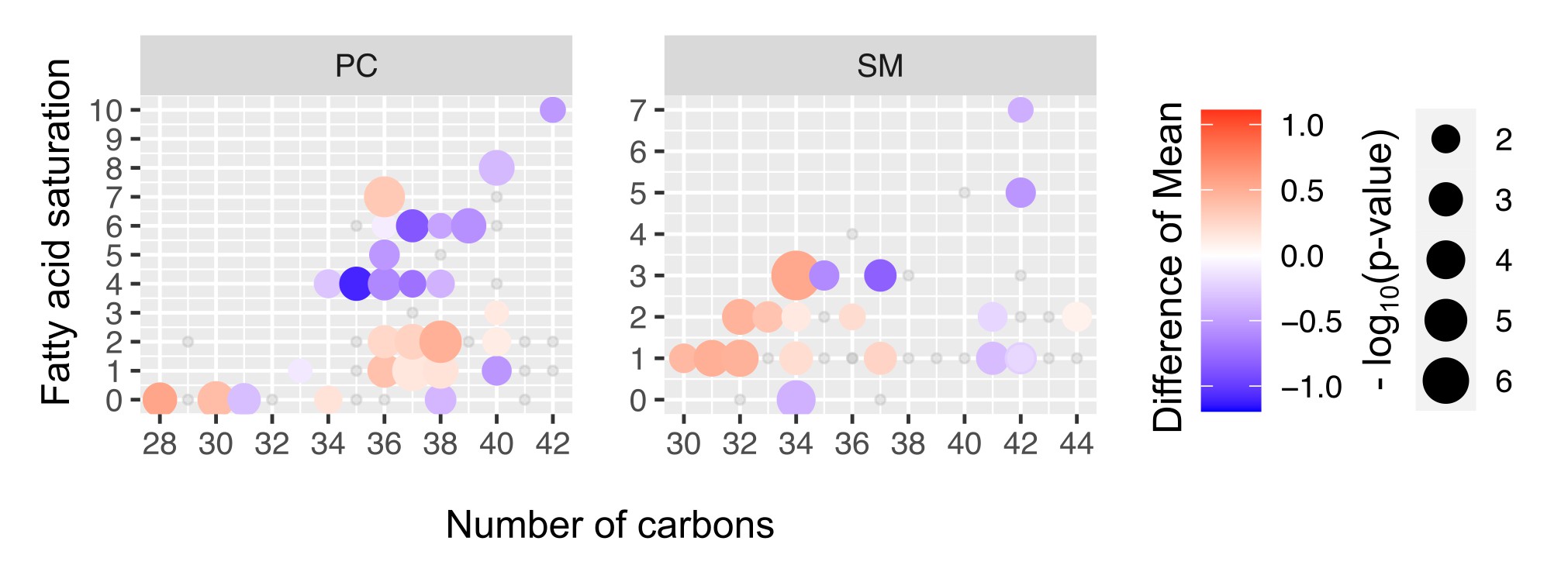
